# The 9-1-1 complex protects ssDNA gaps in BRCA2-deficient cancer

**DOI:** 10.1101/2025.10.07.680950

**Authors:** Helen E. Grimsley, Katherine Courtemanche, Shane Cox, Niamh McDermott, Aman Sharma, Jackson Bright, Jeremy Setton, Mehmet Orman, Simon N. Powell

## Abstract

Single-stranded DNA (ssDNA) gaps are a hallmark of BRCA-deficient cells, yet the mechanisms that safeguard these lesions remain unclear. Through a genome-wide CRISPR screen, we identified the RAD9A-HUS1-RAD1 (9-1-1) complex as essential for the survival of BRCA2-deficient cells through an ATR-independent mechanism. Loss of 9-1-1 in this context leads to the accumulation of PRIMPOL-dependent gaps that fail to undergo post-replicative repair, resulting in pathological expansion and increased DNA damage. This instability is driven by excessive EXO1-mediated degradation, as EXO1 depletion rescues the phenotype. We further demonstrate that the 9-1-1 complex is required for POLζ-dependent gap filling. We propose a model in which ssDNA gaps, when extended beyond a critical length, become inaccessible to TLS-mediated repair and are fully reliant on homologous recombination. These findings establish the 9-1-1 complex as key regulator of ssDNA gap stability and a promising therapeutic target in BRCA2-deficient cancers.

**Graphical Abstract:** 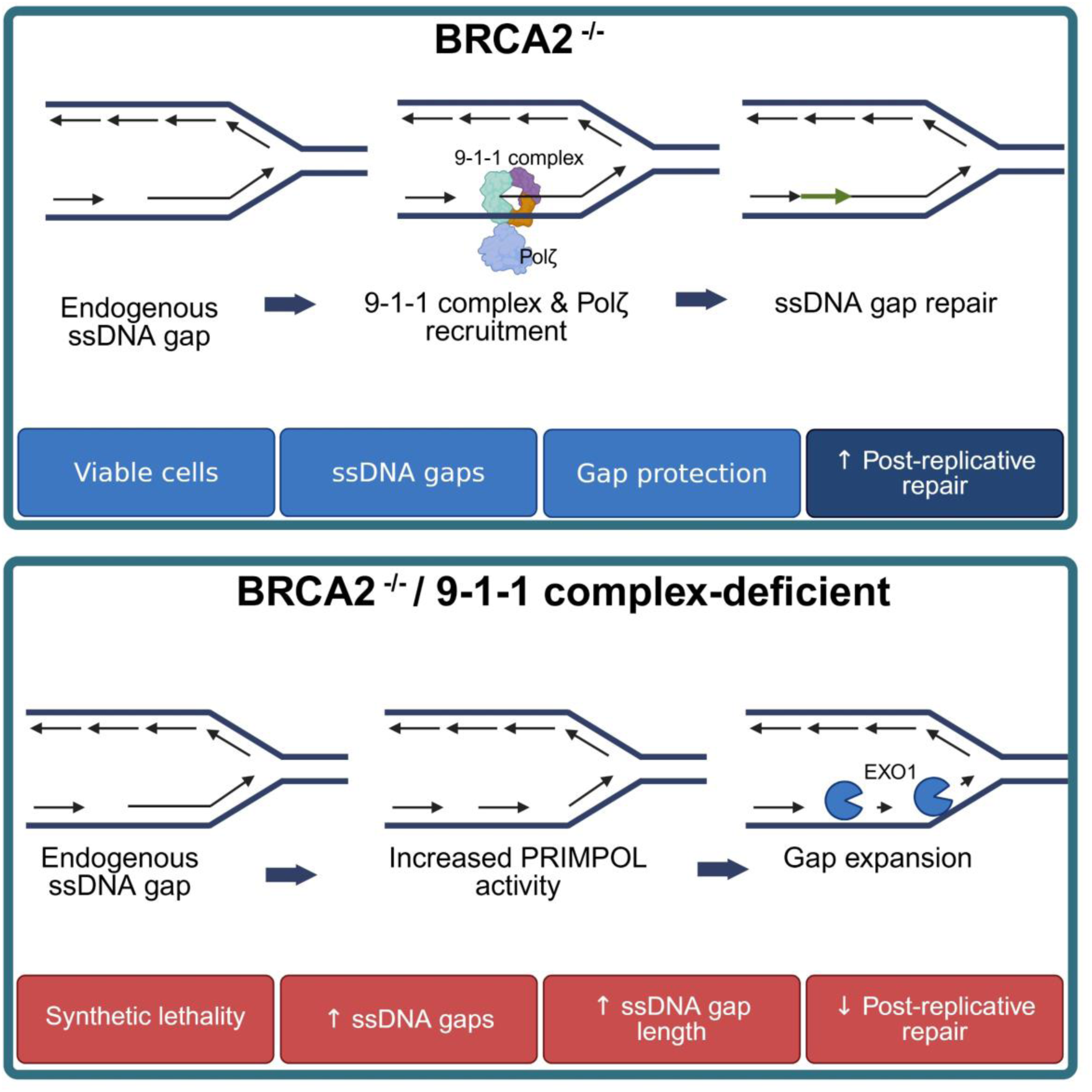

## Introduction

Homologous recombination (HR) is essential for repairing DNA double-strand breaks and resolving replication-associated damage, ensuring accurate genome duplication. BRCA1 and BRCA2 are key mediators of HR, and their loss of function leads to heightened genomic instability, forcing BRCA-deficient cells to rely on alternative DNA repair mechanisms for survival^1^. Recently, single-stranded DNA (ssDNA) gaps have emerged as a significant hallmark of BRCA-deficiency, which contributes to their genomic instability and sensitivity to DNA-damaging drugs, including PARP inhibitors and platinum compounds ^2–8^.

SsDNA gaps arise during DNA replication when lagging-strand synthesis is incomplete or when progression of the leading strand is impeded ^9^. Multiple factors contribute to gap formation, including replication stress (e.g., nucleotide depletion or replication fork stalling), DNA damage (e.g., lesions caused by oxidative stress, UV radiation, or alkylating agents), defects in lagging strand synthesis (e.g., inefficient processing of Okazaki fragments), and impairments in DNA damage tolerance pathways such as template switching, fork reversal, or repriming ^9^. In BRCA-deficient cells, the failure to properly slow or restrain DNA replication in response to genotoxic stress allows DNA synthesis to continue over damaged or compromised templates, promoting the formation of unreplicated regions that manifest as ssDNA gaps ^10^. This can be facilitated by the primase-polymerase PRIMPOL, which allows for repriming downstream of DNA lesions or replication stress, thereby bypassing the damage but leaving unresolved ssDNA regions ^11^. This process is essential for replication progression when fork reversal is not possible ^12^. BRCA-deficient cells also exhibit defects in Okazaki fragment processing and ligation, leading to the accumulation of gaps on the lagging strand ^2^.

Accurate repair and processing of ssDNA gaps are critical for maintaining genome stability and preventing cell death. Multiple proteins act to protect these gaps and limit the accumulation of DNA damage. Current models suggest that eukaryotic cells employ two main post-replicative gap repair strategies: translesion synthesis (TLS) and template switching (TS). TLS relies on the recruitment of low-fidelity DNA polymerases to bypass lesions directly, while TS uses the newly synthesized sister chromatid as a template for gap filling, in a manner analogous to strand invasion during HR, requiring RAD51 function ^13,14^. If left unrepaired, ssDNA gaps can be converted into other DNA lesions, including double-strand breaks, that ultimately require HR for resolution ^15–18^. BRCA1 and BRCA2 play essential roles in recognizing these ssDNA gaps and facilitating their repair. In their absence, ssDNA gaps persist due to inefficient repair and suppression mechanisms.

Paradoxically, BRCA-deficient cells remain viable under unchallenged conditions despite the accumulation of endogenous ssDNA gaps. This vulnerability has been therapeutically exploited by strategies that either promote further ssDNA gap formation or inhibit the compensatory repair pathways these cells depend on for survival ^2–8,19^. While it is clear that these backup pathways are essential, the specific proteins that directly protect ssDNA gaps from expanding into lethal lesions in BRCA2-deficient cells remain largely undefined.

To gain deeper insight into the mechanisms that safeguard ssDNA gaps in BRCA-deficient cells, we conducted a whole-genome CRISPR knockout screen and identified each member of the RAD9A-HUS1-RAD1 (9-1-1) complex, as well as its clamp loader RAD17, as essential in a BRCA2-deficient background. The loss of the 9-1-1 complex, a DNA clamp loader canonically known for its role in activating the ATR checkpoint pathway, was strikingly specific to BRCA2 loss, pointing to a distinct mechanism in gap surveillance.

We demonstrate that the 9-1-1 complex plays a critical ATR-independent role in suppressing the accumulation of PRIMPOL-mediated ssDNA gaps thereby helping to maintain genomic stability in the absence of BRCA2. To understand how the complex exerts this protective effect, we needed to directly visualize gap dynamics. A major challenge in studying gap dynamics has been the direct detection of ssDNA gaps within double-stranded DNA fibers. To address this limitation, we developed Single Molecule Analysis of ssDNA Gaps (SMAss), a modified DNA combing assay that enables quantitative measurement of both gap density (gaps per unit length) and gap length. These parameters have been demonstrated to serve as direct indicators of ssDNA damage severity. Notably, longer and more frequent gaps are preferentially targeted by the nucleases MRE11 and EXO1, which promote their expansion and ultimate progression into double-strand breaks ^16^. For this reason, gap density and length represent strong predictors of overall genomic stability, particularly in BRCA-deficient cells.

SMAss revealed that 9-1-1 loss triggers extensive EXO1-mediated degradation of ssDNA gaps, rendering them incompetent for POLζ-dependent gap filling. We propose a model where the 9-1-1 complex acts as a molecular gatekeeper at ssDNA gaps, physically restraining EXO1 to maintain gaps within a size range accessible to TLS polymerases. When this protection fails, gaps exceed a critical length, becoming irreparable and driving genomic catastrophe.

Our work defines an ATR-independent, scaffolding role for the 9-1-1 complex in ssDNA gap protection, revealing a fundamental mechanism of replication stress tolerance and a new therapeutic vulnerability in BRCA2-deficient cancers. These results establish the 9-1-1 complex as a potential therapeutic target for BRCA2-deficient cancers, highlighting the rationale for developing small-molecule inhibitors to disrupt this pathway. By uncovering key mechanisms governing ssDNA gap stability, our study enhances our understanding of genome maintenance and opens new avenues for novel therapeutic strategies in BRCA2-deficient tumors.

## Results

### Synthetic lethality between BRCA2 loss and the 9-1-1 complex

To identify additional back-up pathways in BRCA2-deficient tumors, we performed a BRCA2 loss-of-function genome-wide CRISPR knockout screen using the Brunello Library with unperturbed isogenic DLD1 WT and DLD1 BRCA2^-/-^ cells. In addition to genes known to be synthetic lethal with BRCA2 loss of function (e.g., POLQ and RAD52), the screen identified all three members of the 9-1-1 complex (RAD9A, RAD1, and HUS1), as well as its clamp loader RAD17 and its binding partner TOPBP1, to be significant hits (Figure 1A, Supplementary File 1). This finding is further supported by additional BRCA2^-/-^ essentiality screens, in which components of the 9-1-1 complex also emerge as significant hits ^20–23^. To confirm our observation, we depleted RAD9A, HUS1, and RAD1 in DLD1 WT and DLD1 BRCA2^-/-^ cells. Here we observed reduced viability with each depletion in the DLD1 BRCA2^-/-^ cells, whilst the DLD1 WT cells remained equally viable relative to the non-targeting control (Figure 1B, Supplementary Figure 1A). Depletion of the 9-1-1 complex was also synthetically lethal in RPE1 BRCA2^-/-^ compared to RPE1 WT, and in H1299 BRCA2 knockout clone, 2D7, compared to H1299 WT (Supplementary Figure 1B-E).

**Figure 1.**
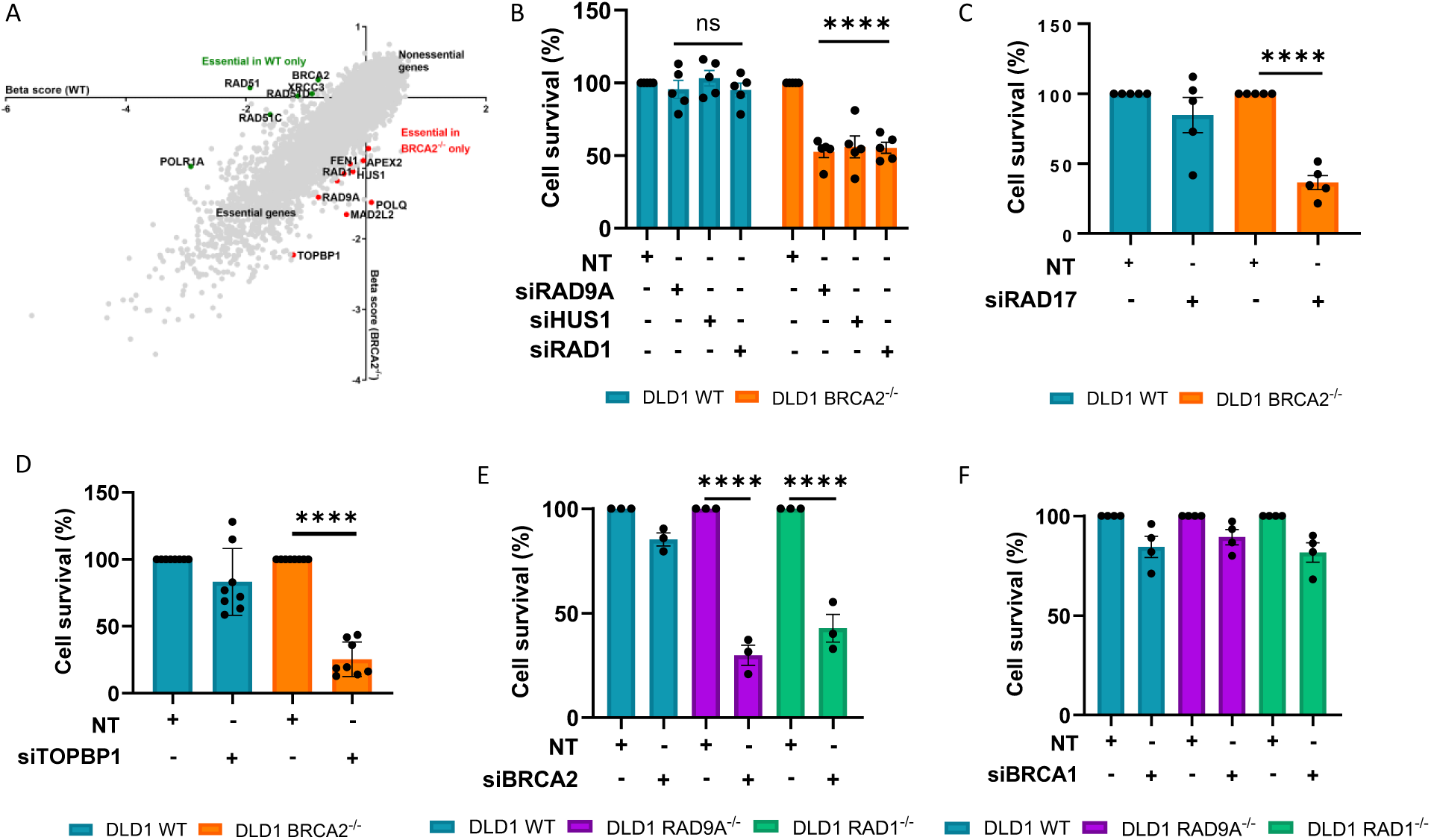
BRCA2 and 9-1-1 complex loss is synthetic lethal. (A) Results of gene essentiality screen in BRCA2-proficient and-deficient DLD1 cells. Plotted is the beta score, with a lower score indicating essentiality. (B-D) Clonogenic assays in the BRCA2-proficient and-deficient DLD1 cells with siRNA depletion of (B) RAD9A, HUS1, RAD1 (C) RAD17 and (D) TOPBP1, n = ≥ 3, mean + SEM, ns= p>0.05, ***p <0.0001, paired t test. (E-F) Clonogenic assays using RAD9A and RAD1-deficient and proficient DLD1 cells with siRNA depletion of (E) BRCA2 and (F) BRCA1, n = ≥ 3, mean + SEM, ns= p>0.05, ****p <0.0001, paired t test. Western blot of lysates shown in Supplementary Figures 1 and 2.

To further validate hits from the CRISPR screen, we next depleted RAD17 and TOPBP1. Synthetic lethality was observed upon depletion of both RAD17 and TOPBP1 in DLD1 BRCA2^-/-^ cells compared to DLD1 WT and in H1299 BRCA2 knockout clone 2D7 compared to H1299 WT (Figure 1C,1D, Supplementary Figure 1F-1G).

To rule out off-target effects of siRNA and enable mechanistic studies of the synthetic lethal interaction, we generated CRISPR-Cas9-mediated RAD9A and RAD1 knockout cell clones in the DLD1 WT cells (DLD1 RAD9A^-/-^, DLD1 RAD1^-/-^). Corroborating the results of the 9-1-1 complex siRNA depletion assays, depletion of BRCA2 in the knockout models further validated the observed synthetic lethal relationship (Figure 1E, Supplementary Figure 2A).

We next sought to understand if synthetic lethality is observed with the combined loss of the 9-1-1 complex and BRCA1. Depletion of BRCA1 mildly affected the clonogenic survival across all cell lines, but there was no significant difference between depletion of BRCA1 in DLD1 RAD9A^-/-^ and DLD1 RAD1^-/-^ compared to DLD1 WT (Figure 1F, Supplementary Figure 2B). Lack of synthetic lethality was also observed with the depletion of the 9-1-1 complex in RPE1 BRCA1^-/-^ cells compared to RPE1 WT (Supplementary Figure 1B).

Our findings, along with others, show that loss of the 9-1-1 complex is synthetic lethal with the loss of BRCA2 across multiple cell line models. This synthetic lethal relationship is also observed through loss of the 9-1-1 complex clamp loader, RAD17, suggesting that the loading of the 9-1-1 complex onto DNA is critical in BRCA2-deficient cells. Our findings demonstrate that this synthetic lethal effect is specific to BRCA2 loss, as no comparable effect was observed with BRCA1 loss. These findings reveal the 9-1-1 complex, and its loader RAD17, as critical for compensatory ssDNA repair in BRCA2-deficient cancer cells, presenting potentially promising therapeutic targets for selective killing of these tumors.

### The 9-1-1 complex promotes post-replicative repair of PRIMPOL-mediated ssDNA gaps in BRCA2-deficient cells

Single-stranded DNA (ssDNA) gaps have emerged as critical contributors to genome instability, particularly in BRCA-deficient cells, where they are numerically more frequent than double-strand damage ^24^. Structural studies have shown that the 9-1-1 complex is recruited to the 5′ ends of DNA at double-stranded/single-stranded junctions, positioning it as a potential sensor or coordinator of repair at these sites ^25,26^. In light of our finding that defective 9-1-1 complex loading is synthetically lethal with BRCA2 loss, we next explored the possibility that the 9-1-1 complex plays a direct role in the recognition or repair of ssDNA gaps.

First, we assessed the impact of loss of the 9-1-1 complex with BRCA2-depletion on replication fork speed. To assess replication fork progression, cells were sequentially pulse-labeled with the thymidine analogs 5-chloro-2’-deoxyuridine (CldU) (short pulse) and IdU (longer pulse), followed by DNA fiber spreading and immunofluorescent staining. The initial CldU pulse serves as a reference to confirm active replication before the IdU pulse, ensuring that only continuously elongating forks are analyzed. The length of the IdU-labeled tracks was then used as a quantitative readout of replication fork speed. Loss of RAD9A or RAD1 alone led to significantly reduced replication fork speed, while combined loss of the 9-1-1 complex and BRCA2 resulted in only minor additional reduction in replication fork speed (Figure 2A). A reduced replication fork speed was also observed in DLD1 WT upon depletion of BRCA2 (Figure 2A). To determine whether the observed replication slowdown was associated with cell cycle arrest, we performed flow cytometry analysis. There was no significant difference in the percentage of cell cycle phases across the cell lines (Supplementary Figure 3A). This suggests that whilst replication fork speed is locally reduced, there is no role of the overall cell cycle time. To test whether the combined loss of the 9-1-1 complex and BRCA2 leads to an accumulation of ssDNA gaps, we depleted BRCA2 in DLD1 WT, DLD1 RAD9A^-/-^, and DLD1 RAD1^-/-^ cells and pulse-labeled them IdU to allow for the visualization and measurement of DNA fibers (Figure 2B). Sequential treatment with S1 nuclease enzymatically nicks dsDNA opposite an ssDNA gap, resulting in shortened DNA fibers and indicating the presence of gaps ^27^. S1 nuclease treatment following BRCA2 depletion caused a reduction in fiber length in DLD1 WT cells, consistent with known ssDNA gap formation (Figure 2B). A significantly greater reduction was observed in DLD1 RAD9A^-/-^ and RAD1^-/-^ cells, suggesting increased gap accumulation in the absence of the 9-1-1 complex (Figure 2B). Significant shortening of fibers was also observed when RAD17 was depleted in DLD1 BRCA2^-/-^ compared to DLD1 WT (Supplementary Figure 3B).

**Figure 2.**
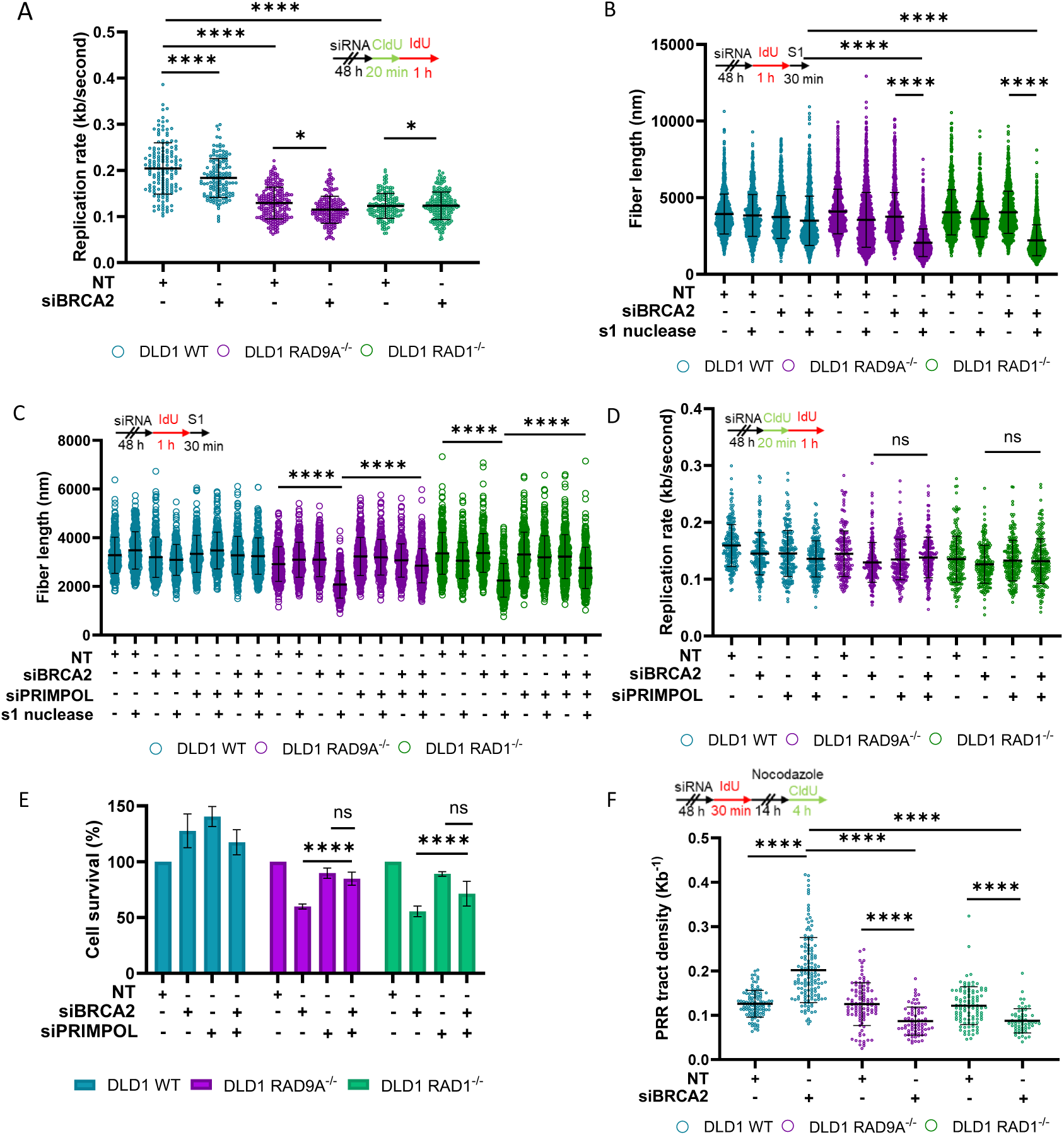
The 9-1-1 loss causes PRIMPOL-mediated gap accumulation and post-replicative repair failure in BRCA2-deficient cells. A) Replication rate with the depletion of BRCA2. Set up of experiment (top) and dot-plot (bottom), n = ≥ 3, at least 100 fibers were analyzed per condition, per replicate, mean + SEM, *p<0.05, ****p <0.0001, paired t test. (B-C) Analysis of DNA fiber length after incubation with and without S1 nuclease. Set up of experiment (top) and dot-plot (bottom) (B) Depletion of BRCA2 and (C) Depletion of BRCA2 and/or PRIMPOL, n = ≥ 3, at least 100 fibers were analyzed per condition, per replicate, mean + SEM, ns= p>0.05, ****p <0.0001, paired t test. (D) Replication rate with the depletion of BRCA2 and/or PRIMPOL. Set up of experiment (top) and dot-plot (bottom), n = ≥ 3, at least 100 fibers were analyzed per condition, per replicate, mean + SEM, *p<0.05, ****p <0.0001, paired t test. (E) Clonogenic survival with co-depletion of BRCA2 and PRIMPOL. (F) Post-replicative repair assay with depletion of BRCA2. Set up of experiment (top) and dot-plot (bottom), n = ≥ 3, at least 20 PRR events were analyzed per condition, per replicate, mean + SEM, ns= p>0.05, ****p <0.0001, paired t test. Western blot of lysates shown in Supplementary Figure 3.

One mechanism by which ssDNA gaps form in BRCA-deficient cells is through PRIMPOL-mediated repriming, which bypasses replication obstacles but at the cost of leaving behind ssDNA gaps behind the fork ^28^. We next investigated whether PRIMPOL activity contributes to the accumulation of ssDNA gaps under the synthetic lethal conditions. Co-depletion of BRCA2 and PRIMPOL in the S1-treated DLD1 RAD9A^-/-^ and DLD1 RAD1^-/-^ cells resulted in longer fiber length compared to BRCA2 depletion alone (Figure 2C, Supplementary 3C). These findings indicate that PRIMPOL contributes to ssDNA gap formation under conditions of combined BRCA2 and 9-1-1 complex loss. Previous work showed that replication forks slow down with fork reversal, but PRIMPOL-mediated repriming does not result in measurable replication fork slowing ^28^. Replication forks were slowed with the loss of the 9-1-1 complex alone; however, no further change in fork speed was observed with the combined loss of the 9-1-1 complex and BRCA2, supporting that the observed ssDNA gaps are PRIMPOL-mediated (Figure 2D). We next sought to determine whether PRIMPOL-mediated gap accumulation contributes to cell death. Clonogenic assay demonstrates that with loss of PRIMPOL, a rescue of the synthetic lethal phenotype was observed (Figure 2E). This suggests that PRIMPOL-generated ssDNA gaps are not only more prevalent but also contribute directly to the synthetic lethality between BRCA2 and the 9-1-1 complex. Together, these data indicate that the ssDNA gaps are generated through PRIMPOL-mediated repriming under the synthetic lethal conditions.

Whilst an increase of PRIMPOL-mediated ssDNA gaps has been observed, we next investigated whether these gaps are being repaired in a post-replicative manner. To test this, we pulse-labeled DLD1 WT, DLD1 RAD9A^-/-^, and DLD1 RAD1^-/-^, BRCA2-depleted cells with IdU to mark DNA synthesized during active replication, resulting in uniform (unifilar) labeling of replication tracks. Immediately afterward, cells were treated with nocodazole for 14 hours to arrest them at the G2/M boundary, providing a window for post-replicative gap-filling. During the final 4 hours of nocodazole treatment, cells were pulse-labeled CldU. By this stage, any CldU incorporation reflects post-replicative DNA synthesis that fills in previously formed ssDNA gaps. These gap-filling repair events appear as discrete CldU patches embedded within the continuous IdU-labeled fibers ^27^. Post-replicative repair (PRR) events are measured in tract density by the ratio of the number of CldU patches to the length of the IdU-labelled DNA fiber ^27^. BRCA2 depletion alone increased the frequency of post-replicative CldU patches, indicating active gap-filling (Figure 2F). However, when combined with 9-1-1 complex loss, this post-replicative repair activity was significantly reduced, suggesting that these gaps persist and are unrepaired (Figure 2F).

### The 9-1-1 complex prevents ssDNA gap expansion and genomic instability

To investigate why the ssDNA gaps fail to undergo post-replicative repair, we sought to understand their dynamics further. We hypothesized that the 9-1-1 complex might act as a protective barrier at the 5’ end of ssDNA gaps, and that its loss could lead to abnormal extension of the ssDNA tract. To test this hypothesis, we developed a new assay to allow for the single-molecule analysis of ssDNA gaps (SMAss) within double-stranded DNA (dsDNA) fibers (Figure 3A). This approach enables direct visualization and quantification of ssDNA gaps at single-molecule resolution, surpassing the limitations of previous indirect and low-resolution DNA fiber assays.

**Figure 3.**
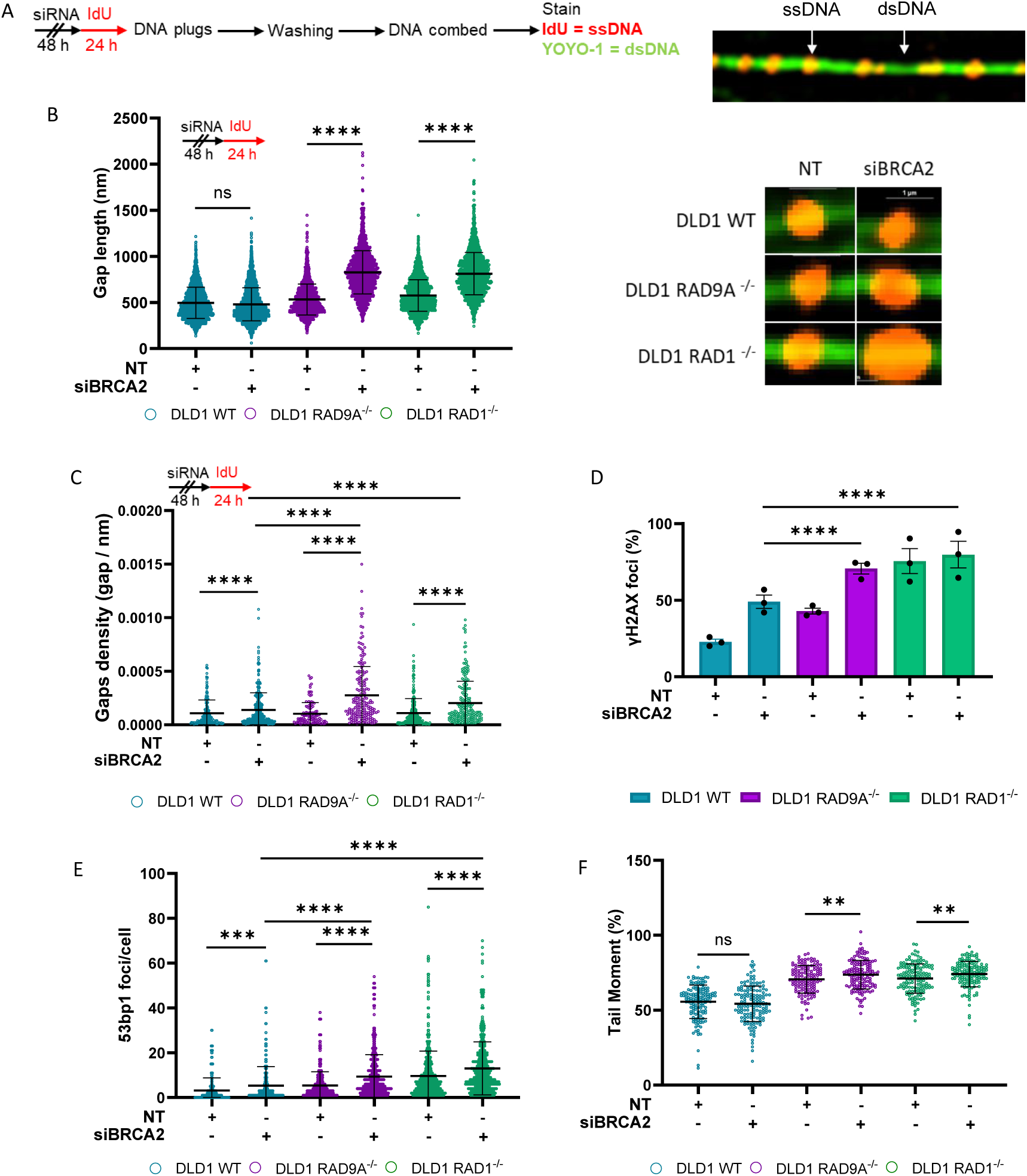
Loss of the 9-1-1 complex results in ssDNA gap expansion and genomic instability. (A) Single Molecule analysis of ssDNA gaps (SMass). Schematic of experiment (left) example of DNA fiber (right). (B) SMass with depletion of BRCA2 with dot-plot (left) and examples of ssDNA gaps for each condition (right), n = 3, at least 100 ssDNA gaps were analyzed per condition, per replicate, mean + SEM, ns= p>0.05, ****p <0.0001, paired t test. (C) Gap density with BRCA2 depletion, set up of experiment (top) dot plot (bottom) n = 3, at least 50 fibers were analyzed per condition, per replicate, mean + SEM, ns= p>0.05, ****p <0.0001, paired t test. (D) Percentage of cells with ≥ 10 γH2AX foci per cell with BRCA2 depletion, n=3, at least 100 cells were analyzed per condition, per replicate mean + SEM, ns= p>0.05, ****p <0.0001, paired t test. (E) Number of 53BP1 foci per cell, n=3, at least 100 cells were analyzed per condition, per replicate, mean + SEM, ns= p>0.05, ****p <0.0001, paired t test. (F) Percentage tail moment from neutral comet assay with BRCA2 depletion, n=3. At least 50 comets were analyzed per condition, per replicate, mean + SEM, ns = non-significant, P** <0.01.

Cells were pulse-labeled with IdU for 24 hours before harvesting for DNA combing under non-denaturing conditions, enabling the detection of ssDNA gaps within dsDNA fibers (Figure 3A). DNA fibers were stained with an anti-BrdU antibody to visualize IdU-labeled ssDNA and counterstained with YOYO-1 dye to specifically mark dsDNA (Figure 3A). Using super-resolution imaging, we were able to detect and quantify ssDNA gaps within the combed DNA fibers (Figure 3A).

Quantitative analysis revealed that the combined loss of the 9-1-1 complex and BRCA2 led to a significant increase in ssDNA gap length compared to BRCA2 loss alone (Figure 3B). Similarly, RAD17 depletion in DLD1 BRCA2^-/-^ cells resulted in longer ssDNA tracts relative to BRCA2 deficiency alone (Supplementary Figure 4A). These findings suggest that the 9-1-1 complex and its loading partner play a critical role in limiting ssDNA gap extension.

Using SMAss, we are also able to calculate the number of ssDNA gaps within each unit length of dsDNA fiber. In confirmation with the observations using the S1 assay, we found that BRCA2 depletion results in a higher number of ssDNA gaps per fiber length (gap density), with a further increase observed under the synthetic lethal conditions (Figure 2B, 3C).

Next, we examined DNA damage markers under the combined deficiency conditions to evaluate the impact of ssDNA gap expansion and impaired post-replicative repair on genome stability. We first assessed γH2AX and 53BP1 foci as indicators of DNA damage (Figure 3D, 3E). Loss of the 9-1-1 complex alone led to an increase in both markers, likely reflecting a deficiency in ATR-dependent replication stress response (Figure 3D, 3E). Notably, BRCA2 depletion further amplified γH2AX and 53BP1 foci, suggesting the activation of a DNA repair response that remains unresolved (Figure 3D, 3E). To corroborate these findings, we used a neutral comet assay to quantify double-strand breaks by measuring the comet tail moment. Loss of the 9-1-1 complex alone increased the tail moment, and this effect was significantly further exacerbated by BRCA2 depletion, indicating elevated genomic instability (Figure 3F).

Collectively, these results demonstrate that disruption of the 9-1-1 complex induces minor signs of genomic instability, which is magnified significantly in the additional absence of BRCA2. These data suggest that decreased cell viability is a result of genomic instability and the secondary production of double-strand breaks acting on the increased number and size of the ssDNA gaps.

### The 9-1-1 complex protects ssDNA gaps from EXO1-mediated degradation

Next, we sought to understand the mechanism that results in ssDNA gap expansion, and whether this was directly linked to the increased genomic instability and loss of cell viability observed with the combined loss of the 9-1-1 complex and BRCA2. In yeast, the 9-1-1 complex has been found to inhibit or promote DNA degradation by exonucleases based on the absence or presence of 53BP1 ^29,30^. We hypothesized that our observations may be result of dysregulated exonuclease activity.

To test this, we performed colony formation assays with co-depletion of BRCA2 and the exonucleases, DNA2, MRE11, and EXO1, in the DLD1 WT, DLD1 RAD9A^-/-^, and DLD1 RAD1^-/-^ cells. Co-depletion of EXO1 and BRCA2 resulted in a complete rescue of the synthetic lethal phenotype in the DLD1 RAD9A^-/-^ and DLD1 RAD1^-/-^ cells (Figure 4A, Supplementary Figure 5A). Consistent with previous observations, no changes in cell vulnerability were seen when DLD1 WT was co-depleted with EXO1 and BRCA2, showing that EXO1 is not essential for the survival of BRCA2-deficient cells (Figure 4A) ^31^. There was no significant difference with the combined loss of BRCA2 and DNA2 or MRE11 in the DLD1 RAD9A^-/-^ and DLD1 RAD1^-/-^ cells compared to BRCA2 loss alone, suggesting that these exonucleases’ activity does not drive the observed synthetic lethality (Supplementary Figure 5B-5D).

**Figure 4.**
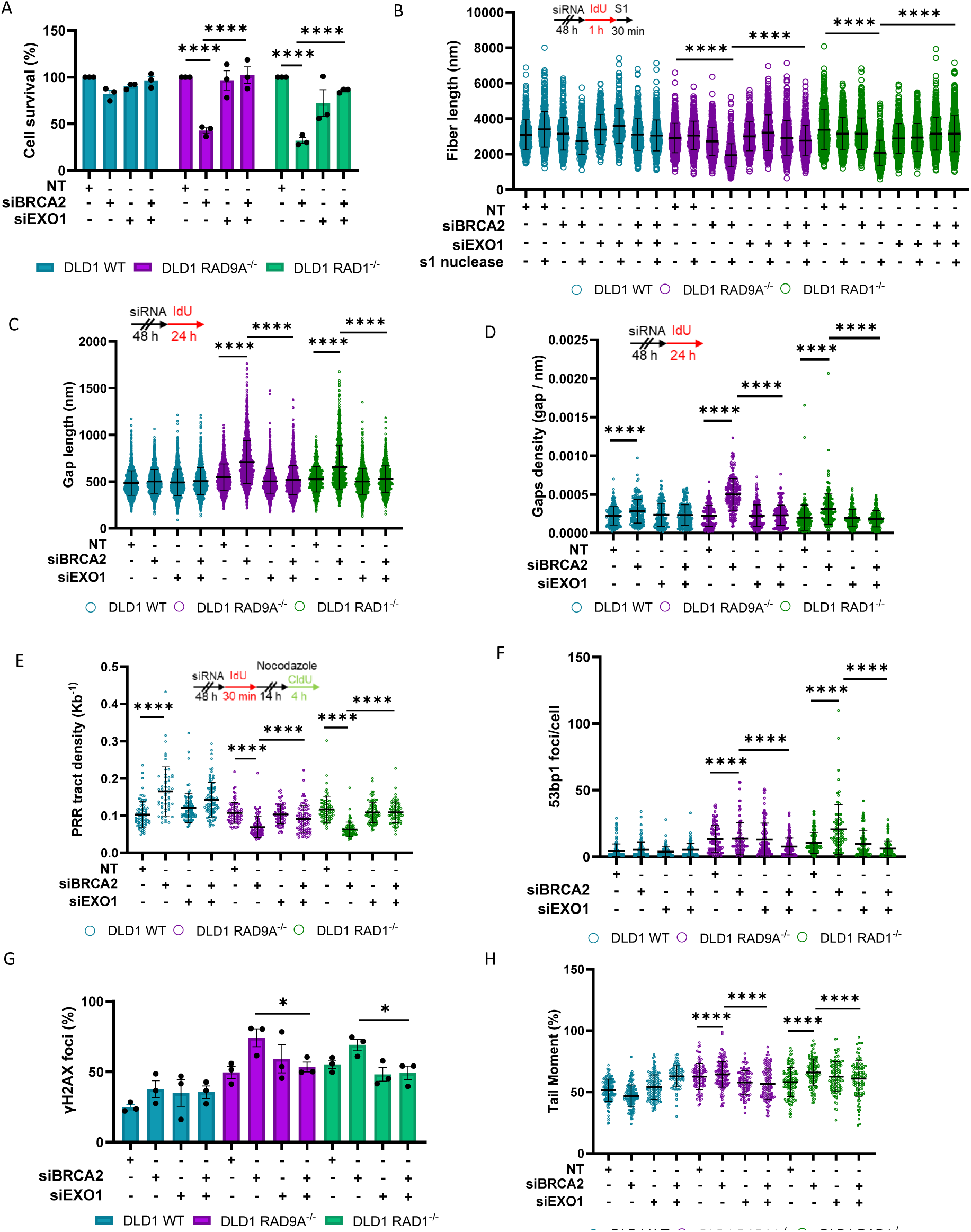
The 9-1-1 complex is required to prevent EXO1 ssDNA gap expansion. (A) Clonogenic assays in the RAD9A or RAD1-proficient and-deficient DLD1 cells with siRNA depletion of BRCA2 and/or EXO1. (B) Analysis of DNA fiber length after incubation with and without S1 nuclease. Set up of experiment (top) and dot-plot (bottom). Depletion of BRCA2 and/or EXO1, n = ≥ 3, at least 100 fibers were analyzed per condition, per replicate, mean + SEM, ns= p>0.05, ****p <0.0001, paired t test. (C) SMass with depletion of BRCA2 and or EXO1. Set up of experiment (top) and dot-plot(bottom). (n = 3, at least 100 ssDNA gaps were analyzed per condition, per replicate, mean + SEM, ns= p>0.05, ****p <0.0001, paired t test). (D) Gap density with BRCA2 and/or EXO1 depletion, set up of experiment (top) dot plot (bottom) n = 3, at least 50 fibers were analyzed per condition, per replicate, mean + SEM, ns= p>0.05, ****p <0.0001, paired t test. (E) Post replicative repair assay with depletion of BRCA2 and/or EXO1. Set up of experiment (top) and dot-plot (bottom) (n = ≥ 3, at least 20 PRR events were analyzed per condition, per replicate, mean + SEM, ns= p>0.05, ****p <0.0001, paired t test). (F) Number of 53BP1 foci per cell (n=3, at least 100 cells were analyzed per condition, per replicate, mean + SEM, ns= p>0.05, ****p <0.0001, paired t test). (G) Percentage of cells with ≥ 10 γH2AX foci per cell with BRCA2 depletion, n=3, at least 100 cells were analyzed per condition, per replicate mean + SEM. (H) Percentage tail moment from neutral comet assay with BRCA2 depletion, n=3. At least 50 comets were analyzed per condition, per replicate, mean + SEM, ns = non-significant, P** <0.01.

Previous studies have shown that EXO1, MRE11, and DNA2 can act at ssDNA gaps, prompting us to evaluate whether co-depletion of BRCA2 with the aforementioned exonucleases influences gap accumulation ^16,32,33^. Co-depletion of BRCA2 and EXO1 resulted in longer DNA fibers in DLD1 RAD9A^-/-^ and DLD1 RAD1^-/-^ cells compared to BRCA2 depletion alone (Figure 4B). This recovery was not observed with the co-depletion of BRCA2 and EXO1 in DLD1 WT (Figure 4B). A modest rescue effect was observed with the co-depletion of MRE11 or DNA2 under the synthetic lethal conditions, although it was not as pronounced as that seen with co-depletion of BRCA2 and EXO1 (Supplementary Figure 5E, 5F).

Taken together, these findings suggest that while MRE11 and DNA2 may contribute to ssDNA gap processing, their activity is not the primary driver of the synthetic lethality observed upon combined loss of the 9-1-1 complex and BRCA2. We therefore directed our subsequent analyses toward EXO1, whose depletion produced the most substantial rescue effect.

To confirm that the ssDNA gaps rescued by EXO1 were those generated by the loss of the 9-1-1 complex, we induced ssDNA gaps in the DLD1 WT and DLD1 BRCA2^-/-^ cells through treatment with the PARP inhibitor Olaparib. Shorter DNA fibers were observed when the DLD1 BRCA2^-/-^ cells were treated with Olaparib as previously established (Supplementary Figure 6A)^2^. However, there was no rescue of fiber length with the depletion of EXO1, indicating that these gaps are distinct from those observed in the synthetic lethal relationship (Supplementary Figure 6A).

We next performed SMAss to measure the length and density of the ssDNA gaps. We found that co-depletion of EXO1 and BRCA2 in the DLD1 RAD9A^-/-^ and DLD1 RAD1^-/-^ cells reduced the gap length compared to BRCA2 depletion alone, suggesting that gap extension is EXO1-dependent (Figure 4C). In confirmation with the S1 assay, we also found a rescue in gap density (Figure 4B, 4D).

To understand if depletion of EXO1 under the synthetic lethal conditions can rescue ssDNA gap repair, we assessed the post-replicative repair of ssDNA gaps. Here, we observed a rescue in post-replicative events when DLD1 RAD9A^-/-^ and DLD1 RAD1^-/-^ cells were co-depleted with BRCA2 and EXO1 compared to BRCA2 depletion alone (Figure 4E). This indicates that gap expansion by EXO1 results in reduced post-replicative DNA repair. However, it was noted that whilst the depletion of EXO1 resulted in a rescue to that of DLD1 RAD9A^-/-^ and DLD1 RAD1^-/-^ non-target PRR tract density, it did not replicate the observation of BRCA2 or BRCA2/EXO1 depletion in DLD1 WT (Figure 4E). These findings indicate that EXO1 depletion only partially restores ssDNA gap repair, suggesting that the 9-1-1 complex carries out additional, EXO1-independent functions at these lesions.

Finally, we evaluated whether loss of EXO1 can reduce the pronounced DNA damage markers observed in the synthetic lethal conditions. We detected a reduction in both of γH2AX and 53BP1 foci with loss of EXO1 and BRCA2 in DLD1 RAD9A^-/-^ and DLD1 RAD1^-/-^ cells compared to BRCA2 depleted DLD1 RAD9A^-/-^ and DLD1 RAD1^-/-^ (Figure 4F, 4G). A reduction of DNA damage was observed when assessing the tail moment with co-depleted EXO1 and BRCA2 in the DLD1 RAD9A^-/-^ and DLD1 RAD1^-/-^ cells compared to BRCA2-depleted DLD1 RAD9A^-/-^ and DLD1 RAD1^-/-^ (Figure 4H). Together, these data suggest that ssDNA gap expansion by EXO1 results in greatly increased genomic instability, driving the synthetic lethal phenotype.

### The 9-1-1 complex facilitates translesion synthesis at ssDNA gaps

Translesion synthesis (TLS) proteins are essential for the repair of ssDNA gaps, especially in BRCA-deficient cells, where canonical repair pathways are compromised ^7,8,34,35^. Emerging evidence suggests that the 9-1-1 complex may play a supportive role in facilitating TLS ^36–38^. Of note, in yeast studies, the 9-1-1 complex has been shown to interact with POLζ, a B-family DNA polymerase central to TLS activity. Furthermore, POLζ is essential in the gap-filling of PRIMPOL-mediated ssDNA gaps ^8^.

Recently, Predictome—an AlphaFold-Multimer-based platform for protein interaction prediction—has demonstrated strong experimental validation ^39^. Using this tool, we identified RAD9A as a top predicted interactor of REV3L and REV1, with 149 and 388 contact points, respectively (Figure 5A, 5B, Supplementary Figure 7A, 7B). REV3L serves as the catalytic subunit of POLζ, while REV1 is critical for its recruitment ^40^. High-confidence structural metrics and omics-informed classifier scores support these predictions.

**Figure 5.**
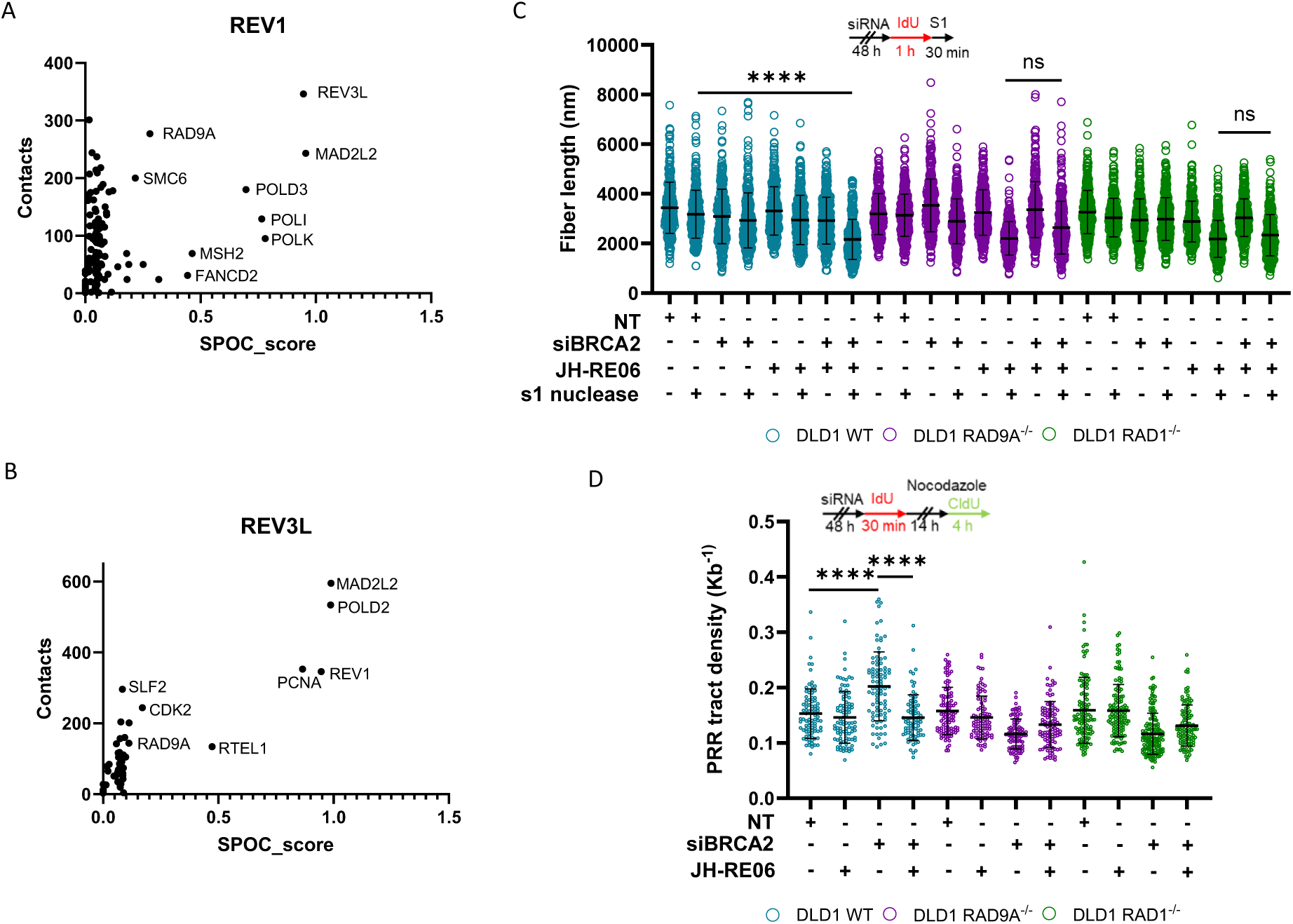
The 9-1-1 complex promotes POLζ-mediated gap filling. (A-B) Data selected from Predictome—an AlphaFold-Multimer-based platform for protein interaction prediction. Plotted is the number of predicted contacts against SPOC score (The structure prediction and omics-informed classifier score, which considers structural and biological features of each contact pair). Shown are predictive contacts for (A) REV1 and (B) REV3L. (C) Analysis of DNA fiber length after incubation with and without S1 nuclease. Set up of experiment (top) and dot-plot (bottom). Depletion of BRCA2 with/without JH-RE06 treatment, n = ≥ 3, at least 100 fibers were analyzed per condition, per replicate, mean + SEM, ns= p>0.05, ****p <0.0001, paired t test. (D) Post replicative repair assay with depletion of BRCA2 with/without JH-RE06 treatment. Set up of experiment (top) and dot-plot (bottom), n = ≥ 3, at least 20 PRR events were analyzed per condition, per replicate, mean + SEM, ns= p>0.05, ****p <0.0001, paired t test.

Previous studies have shown that the small molecule JH-RE-06 disrupts the error-prone gap-filling pathway necessary for resolving ssDNA gaps that arise during replication repriming by PRIMPOL^8^. In BRCA-deficient cells, this exacerbates the accumulation of ssDNA ^8^. Mechanistically, JH-RE-06 inhibits the interaction between REV1 and REV7, key subunits of POLζ, by inducing REV1 dimerization, thereby blocking POLζ recruitment ^41^.

To determine whether the 9-1-1 complex collaborates with POLζ at ssDNA gaps, we treated DLD1 WT, DLD1 RAD9A^-/-^, and DLD1 RAD1^-/-^ cells with JH-RE-06 and assessed its impact on ssDNA gaps. Using the S1 nuclease assay, inhibition of REV1 with JH-RE-06 significantly reduced DNA fiber length in 9-1-1 complex–deficient cells; however, additional depletion of BRCA2 in these cells did not further reduce fiber length (Figure 5C). In contrast, in DLD1 WT cells, BRCA2 depletion combined with JH-RE-06 treatment led to clear DNA fiber shortening (Figure 5C). This suggests no additional effect is observed with REV1 inhibition with inactivation of the 9-1-1 complex.

We next examined the effect of JH-RE-06 on post-replicative repair. Inhibition of REV1 did not rescue the PRR defect in BRCA2-depleted DLD1 RAD9A^-/-^ and RAD1^-/-^ cells (Figure 5D).

Interestingly, in DLD1 WT cells, BRCA2 depletion alone led to increased PRR tract density, but this was significantly reduced to baseline levels with JH-RE-06 treatment, suggesting the requirement of REV1 for post-replicative repair (Figure 5D).

This lack of additive effect indicates that REV1 and the 9-1-1 complex operate in the same repair pathway, supporting an epistatic relationship. These findings suggest that the 9-1-1 complex promotes effective ssDNA gap filling through a POLζ-dependent mechanism.

### Loss of 9-1-1 complex and BRCA2 synthetic lethality is ATR-independent

The 9-1-1 complex is known to activate DNA damage checkpoint signaling of ATR and CHK1 in response to replication stress ^42^. The 9-1-1 complex facilitates the recruitment and activation of ATR and TOPBP1 at sites of replication stress. Our CRISPR-Cas9 viability screen results indicated that ATR and ATRIP were not significant hits of synthetic lethality with BRCA2 loss, with low beta values of-0.34952 and-0.34763, respectively (Supplementary File 1). This led us to hypothesize that the 9-1-1 complex has a role in cell viability independent of the ATR-signaling checkpoint in BRCA2-deficient cells. We assessed the impact of RAD9A and RAD1 loss on downstream ATR and CHK1 signaling (Supplementary Figure 8). Upon hydroxyurea treatment, we observed a substantial increase in phosphorylation at T1989 in ATR and S435 in CHK1 in DLD1 WT (Supplementary Figure 8). However, with the loss of the 9-1-1 complex, we observed weaker phosphorylation of ATR and CHK1 (Supplementary Figure 8). These findings suggest that despite the loss of the 9-1-1 complex, ATR activation is still viable, albeit at a reduced level compared to DLD1 WT.

We next assessed the impact of ATR inhibition (ATRi; using VE-822) on cell survival in the context of 9-1-1 complex loss and BRCA2 depletion. We observed a reduction in cell survival with ATRi treatment in DLD1 WT, DLD1 RAD9A^-/-^, and DLD1 RAD1^-/-^ (Figure 6D). A further reduction in cell survival was seen upon depletion of BRCA2 in all cell lines. Notably, the relative sensitivity to ATR inhibition remains consistent across all conditions, indicating that the synthetic lethality between BRCA2 deficiency and 9-1-1 loss cannot be attributed to compromised ATR signaling. This supports an ATR-independent function for the 9-1-1 complex (Figure 6A).

**Figure 6.**
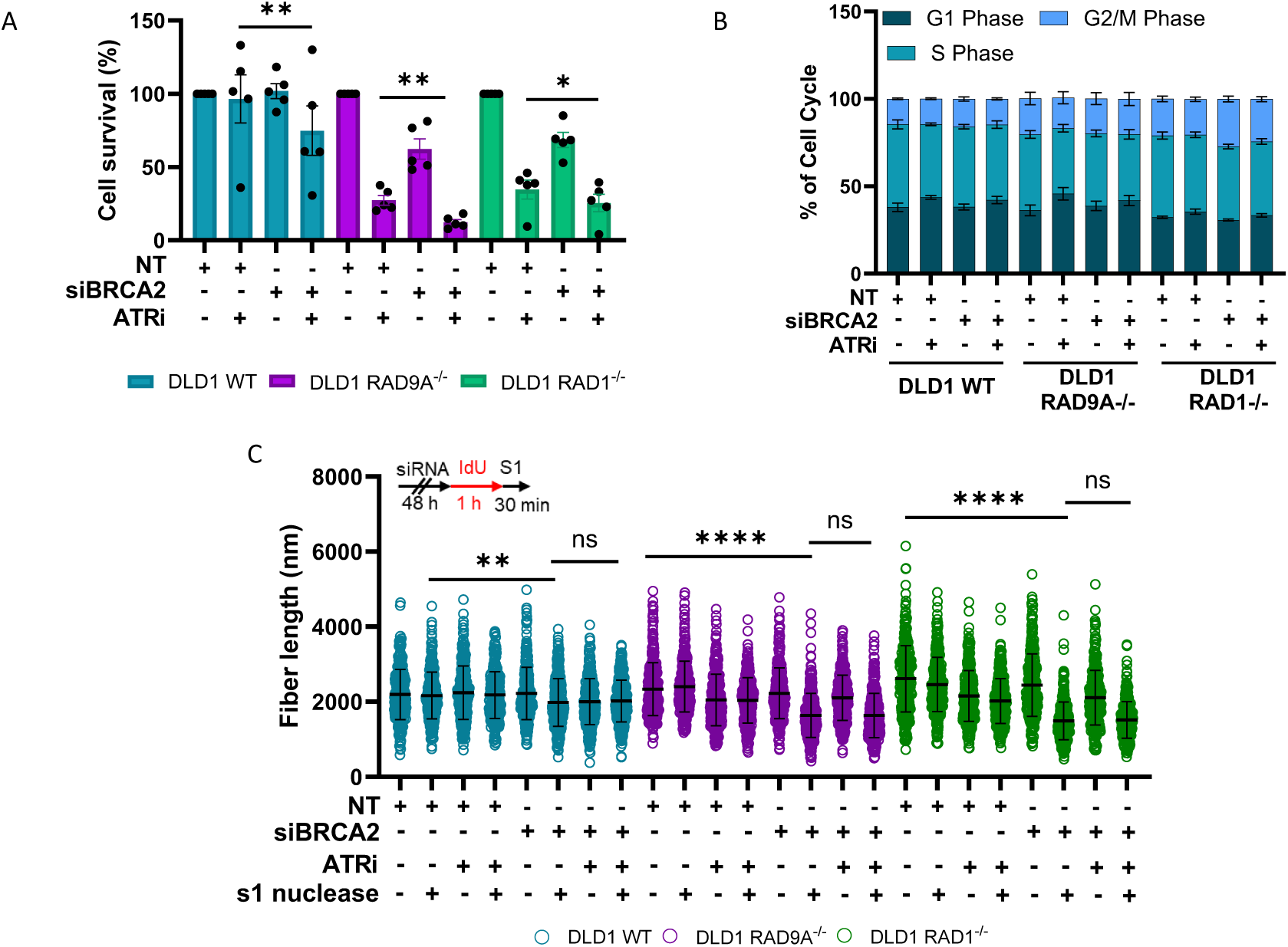
The synthetic lethal relationship is ATR-independent. (A) Clonogenic assays using RAD9A and RAD1-deficient and proficient DLD1 cells with siRNA depletion of BRCA2 and treatment with ATR inhibitor n = ≥ 3, mean + SEM, *p<0.05, **p <0.01, paired t test. (B) Cell cycle analysis of RAD9A/RAD1-deficient and proficient cell lines with BRCA2 depletion and treated with and without an ATR inhibitor, n=3, mean + SEM. (C) Analysis of DNA fiber length after incubation with and without S1 nuclease. Set up of experiment (top) and dot-plot (bottom). Depletion of BRCA2 with/without ATR inhibitor treatment, n = ≥ 3, at least 100 fibers were analyzed per condition, per replicate, mean + SEM, ns= p>0.05, ****p <0.0001, paired t test.

We next considered the impact of ATR inhibition and loss of BRCA2 on the cell cycle in the DLD1 WT, RAD9A^-/-^ and RAD1^-/-^ cell lines. No significant difference was observed in the cell cycle profiles with loss of BRCA2 alone (Figure 6B). However, ATR inhibition resulted in a small increase in the percentage of cells in G1 phase, and a reduced number of cells in S phase in each of the cell lines (Figure 6B). Notably, there was no significant difference between combined BRCA2 loss and ATR inhibition compared to ATR inhibition alone, suggesting that there is no role of the overall cell cycle time or global cell cycle checkpoints in the local DNA damage response (Figure 6B).

Finally, we investigated whether ATR loss influences the accumulation of ssDNA gaps in BRCA2-deficient cells. In contrast to loss of the 9-1-1 complex, combined BRCA2 deficiency and ATR inhibition did not induce detectable ssDNA gap accumulation (Figure 6C). Furthermore, ATR inhibition failed to further enhance gap formation under conditions of synthetic lethality (Figure 6C). Collectively, these findings support the conclusion that the 9-1-1 complex exerts an ATR-independent function in maintaining cell viability in the context of BRCA2-deficiency.

## Discussion

Single-stranded DNA (ssDNA) gaps have emerged as a hallmark of BRCA1/BRCA2-deficiency, driving at least part of the sensitivity to chemotherapeutics such as PARP inhibitors and platinum compounds ^2–4,4,6–8,24,43^. How these ssDNA gaps are safeguarded to allow for continued cell viability whilst unchallenged by therapeutic agents is not understood. Our study redefines the essential role of the 9-1-1 complex in cancer cell survival, uncoupling it from its canonical ATR-checkpoint function and establishing it as a direct protector of ssDNA gaps. We demonstrate that in BRCA2-deficient cells, the 9-1-1 complex functions as a molecular gatekeeper that restrains EXO1-mediated gap expansion and facilitates POLζ-dependent gap filling, thereby maintaining gaps within a tolerable, repairable state.

### The 9-1-1 Complex as a BRCA2-Specific Survival Factor

Our work identifies a previously unrecognized synthetic lethal interaction between BRCA2 loss and 9-1-1 complex disruption. Along with the 9-1-1 complex, we identified RAD17 to be essential in a BRCA2-deficient background, suggesting that the loading of the 9-1-1 complex onto the DNA is necessary in BRCA2-deficient cells (Figure 1). Unexpectedly, the synthetic lethal interaction appears to be specific to BRCA2-deficiency, as BRCA1-deficient cells tolerate loss of the 9-1-1 complex (Figure 1F, Supplementary Figure 1B). This observation aligns with previous findings, including the Durocher lab’s BRCA1 synthetic lethality screen, in which components of the 9-1-1 complex were not identified as significant hits ^21^.

This specificity reveals fundamental differences in how BRCA1 and BRCA2 regulate ssDNA gap metabolism. The most compelling explanation centers on the opposing roles of these factors in regulating EXO1-mediated nucleolytic processing. BRCA1, in complex with BARD1, directly binds and activates EXO1 ^44^. In the absence of BRCA1, EXO1 may not be efficiently recruited or active, making its nucleolytic degradation severely compromised, reducing the nucleolytic pressure on ssDNA gaps and thereby diminishing the protective function of the 9-1-1 complex. If EXO1 activity is already limited in BRCA1-deficient backgrounds, the additional loss of 9-1-1-mediated protection may not create sufficient gap expansion to trigger synthetic lethality. However, alternative explanations warrant consideration, with further investigation of EXO1 recruitment dynamics and chromatin modifications in a BRCA1 and BRCA2 background will help clarify the molecular basis for this specificity. This work, along with others, underscores the importance of dissecting the distinct roles of BRCA1 and BRCA2 in ssDNA gap metabolism^24^.

### 9-1-1 Complex Function in Gap Protection and Repair

Structural studies have demonstrated that the 9-1-1 complex is recruited to the 5’ end of double-stranded/single-stranded DNA (dsDNA/ssDNA) junctions, including both recessed ends and ssDNA gaps ^25,26,45^. In this study, we reveal that loss of the 9-1-1 complex and its clamp loader, RAD17, leads to the accumulation and lengthening of ssDNA gaps (Figure 2A, 3B, 3C Supplementary 3A, 4A). Our findings suggest that these gaps predominantly arise through PRIMPOL-mediated repriming, as both their accumulation and associated cell viability defects are rescued by PRIMPOL depletion (Figure 2B, 2D).

A central finding is that the 9-1-1 complex restricts the expansion of ssDNA gaps. To directly assess gap extension, we developed a high-resolution DNA combing technique, Single Molecule analysis of ssDNA Gaps (SMAss), capable of measuring ssDNA gap lengths within double-stranded DNA. Under synthetic lethal conditions, we observed a marked increase in ssDNA tract length, indicating extensive gap expansion. In addition to measuring gap length, SMAss enables quantification of ssDNA gap density with greater sensitivity than the widely used S1 nuclease assay, providing more precise and quantitative assessment of gap formation.

Using a modified DNA fiber assay, we further show that under synthetic lethal conditions, these ssDNA gaps persist and are not repaired post-replicatively, whilst BRCA2 loss alone demonstrates an increase in post-replicative repair (PRR) (Figure 2E). It has been previously shown that persistent gaps can expose ssDNA to nucleases or secondary structures, which can ultimately result in point mutations or double-strand breaks ^16,33,46^. The absence of a PRR defect in 9-1-1–deficient cells suggests that the complex is not universally required for PRR but instead becomes essential under conditions of elevated replication stress or when HR is compromised. PRIMPOL-dependent gaps are found almost exclusively in HR-deficient cells. This context-specific requirement underscores its role as a tolerance factor for pathological ssDNA gap structures.

### Molecular Mechanisms of Gap protection

Mechanistically, we identified EXO1 as the key nuclease responsible for ssDNA gap expansion under synthetic lethal conditions (Figure 4). EXO1 resects the 5’-terminated strand of duplex DNA in a 5’ to 3’ direction and functions independently of a helicase partner ^16,47–49^. In our system, EXO1 depletion effectively rescues gap accumulation and extension, partially rescues PRR, and enhances clonogenic survival—supporting a model in which the 9-1-1 clamp acts as a barrier or negative regulator of EXO1 activity at sites of ssDNA gaps (Figure 4A-4E). Consistent with previous findings, EXO1 is not essential for the survival of BRCA2-deficient cells (Figure 4A) ^31^. Notably, depletion of other nucleases involved in long-and short-range resection, DNA2 and MRE11, failed to confer similar clonogenic rescue, highlighting the specific role of EXO1 in this context (Supplementary Figure 5B).

The precise molecular mechanism by which the 9-1-1 complex restricts EXO1-mediated ssDNA gap expansion remains to be fully elucidated. Given that both factors recognize 5’ DNA ends, several models warrant consideration. The 9-1-1 complex may directly compete with EXO1 for binding sites at the 5’ terminus of ssDNA gaps, forming a stable nucleoprotein complex that sterically blocks nuclease access. Alternatively, 9-1-1 loading could induce local chromatin modifications or recruit additional factors that allosterically inhibit EXO1 activity. The ring-shaped architecture of the 9-1-1 clamp suggests it may encircle the DNA backbone, creating a physical barrier that restricts EXO1 processivity beyond a certain distance from the gap origin. Future structural and biochemical studies examining the competitive dynamics between these factors, particularly using single-molecule approaches, will be essential to distinguish between these possibilities and define the kinetic parameters governing gap processing.

The partial restoration of PRR following EXO1 depletion suggests that the 9-1-1 complex contributes to PRR through mechanisms beyond merely restricting EXO1-mediated gap expansion. Notably, prior studies have shown that controlled gap extension by EXO1—and, to a lesser extent, MRE11—is required to facilitate efficient PRR ^33^. Thus, in the absence of sufficient gap expansion, the capacity for effective PRR may be inherently limited.

### Coordination with Translesion synthesis

In addition to its role in limiting gap degradation, the 9-1-1 complex appears to actively promote translesion synthesis (TLS)-mediated repair of ssDNA gaps, likely through both physical and functional interactions with TLS polymerases. Predictome-based network analysis revealed strong predicted interactions between RAD9A and two key TLS components: REV3L, the catalytic subunit of polymerase ζ (POLζ), and REV1, which is essential for POLζ recruitment (Figure 5A, 5B). These predictions are supported by previous studies confirming direct interactions between the 9-1-1 complex and POLζ ^37^.

REV1-Polζ has been shown to facilitate error-prone gap filling of ssDNA gaps generated by PRIMPOL-mediated repriming downstream of DNA lesions, thereby preventing lethal replication fork collapse ^7,8^. In BRCA1/BRCA2-deficient cells, where HR is impaired, survival becomes increasingly dependent on REV1-Polζ activity ^7,8^. In this study, pharmacological inhibition of REV1-Polζ did not further exacerbate the phenotype caused by loss of the 9-1-1 complex, while BRCA2 depletion alone in DLD1 wild-type cells resulted in increased ssDNA accumulation and impaired PRR (Figure 5C, 5D). This suggests an epistatic relationship between REV1-Polζ and the 9-1-1 complex, implicating a shared functional pathway.

Our findings raise important questions about the temporal coordination between 9-1-1-mediated gap protection and POLζ-dependent gap filling. The epistatic relationship we observe between 9-1-1 loss and REV1 inhibition suggests these factors operate within the same pathway, but the precise sequence of events remains unclear. One model we propose is that 9-1-1 loading immediately follows gap formation, creating a protected environment that facilitates subsequent recruitment of REV1-POLζ complexes. This would ensure that TLS machinery gains access to gaps before excessive nucleolytic processing renders them irreparable. Alternatively, 9-1-1 and POLζ may be recruited simultaneously through shared protein interactions or chromatin modifications. The observation that 9-1-1 loss impairs post-replicative repair even when EXO1 is depleted suggests additional functions beyond nuclease restriction, possibly including direct facilitation of TLS polymerase recruitment or maintenance of optimal chromatin structure for repair synthesis. Time-course experiments tracking the sequential recruitment of these factors to individual gaps will be necessary to resolve this temporal relationship. Collectively, these findings support a model in which the 9-1-1 complex is necessary for efficient ssDNA gap filling via a POLζ-dependent mechanism, though whether it facilitates POLζ recruitment, activation, or both at sites of replication stress remains to be determined.

Previous studies have implicated the 9-1-1 complex in coordinating with TLS polymerases including POLQ, which plays a pivotal role in repairing ssDNA gaps in BRCA-deficient cells and its loss is synthetic lethal ^4–6,22,50,51^. Along with RHINO—an interacting partner of the 9-1-1 complex—the 9-1-1 complex has been shown to recruit POLQ-mediated microhomology - mediated end joining (MMEJ) to repair DNA breaks during mitosis, raising the possibility that the observed cell death in this study could be driven by mitotic MMEJ activity ^22^. However, the observation that PRIMPOL depletion rescues the synthetic lethal phenotype suggests that the DNA gaps responsible are primarily PRIMPOL-mediated, thereby distinguishing them from MMEJ-associated lesions and implicating an alternative mechanism of cell death (Figure 2D). Whether the 9-1-1 complex serves as a direct physical recruiter of TLS polymerases or instead modulates the chromatin landscape to facilitate their access remains to be determined.

### ATR-Independent Function

The 9-1-1 complex plays an important role in higher eukaryotes due to its ATR-dependent checkpoint signaling. However, in this study we show that the synthetic lethality observed between BRCA2 and the 9-1-1 complex is independent of canonical ATR checkpoint signaling. While 9-1-1 complex loss impairs ATR activation (as evidenced by reduced CHK1 phosphorylation upon HU treatment), neither ATR nor ATRIP emerged as hits in our CRISPR screen, and pharmacological ATR inhibition only modestly enhanced BRCA2-deficient cell death (Figure 6A, Supplementary Figure 8, Supplementary File 1). Importantly, ATR inhibition with BRCA2 depletion did not recapitulate the synthetic lethal conditions observed with gap accumulation in the S1 nuclease assay (Figure 6C). Previous studies suggest that the 9-1-1 complex functions as a molecular scaffold for DNA repair factors independently of ATR activation ^26,36,42,45^. Building on this, our findings indicate that the 9-1-1 complex plays an ATR-independent role in stabilizing ssDNA gaps, highlighting a distinct function in maintaining replication integrity. This ATR-independent activity highlights a broader principle: sliding clamps may exert essential functions by scaffolding local DNA metabolism independent of downstream kinase cascades. From a therapeutic perspective, this opens the possibility that targeting 9-1-1 could complement ATR inhibition rather than act redundantly, expanding the range of vulnerabilities in BRCA2-mutant tumors.

### Critical Gap Length Threshold

Based on our findings, we propose a model in which the 9-1-1 complex facilitates the timely recruitment of TLS polymerases to ssDNA gaps and prevents their pathological extension (Figure 7). If these gaps are allowed to persist and elongate—particularly due to unchecked EXO1 activity—they may exceed the processing capacity of TLS polymerases and become dependent on HR for repair. In BRCA2-deficient cells, where HR is already impaired, this shift renders the gaps toxic and potentially lethal. This work enhances our understanding of how BRCA2-deficient cells survive despite replication stress and ssDNA gap formation, pointing to the 9-1-1 complex as a key mediator of gap tolerance. However, the molecular basis for this threshold remains undefined. Several factors may contribute to the transition from TLS-competent to TLS-refractory gaps. Longer gaps may adopt secondary structures or protein-DNA complexes that inhibit polymerase loading or processivity. Additionally, extensive nucleolytic processing might remove essential protein landmarks or modify chromatin architecture in ways that favor HR over TLS. The gap length at which this transition occurs likely varies between cell types and depends on the local concentration and activity of repair factors. Quantitative analysis of gap length distributions in our SMAss assay suggests this threshold may lie between 100-500 nucleotides, but precise determination will require controlled gap generation systems and real-time monitoring of repair pathway engagement. Understanding this threshold has important implications for therapeutic strategies, as interventions that push gaps beyond the TLS-accessible range could selectively target HR-deficient cells.

**Figure 7.**
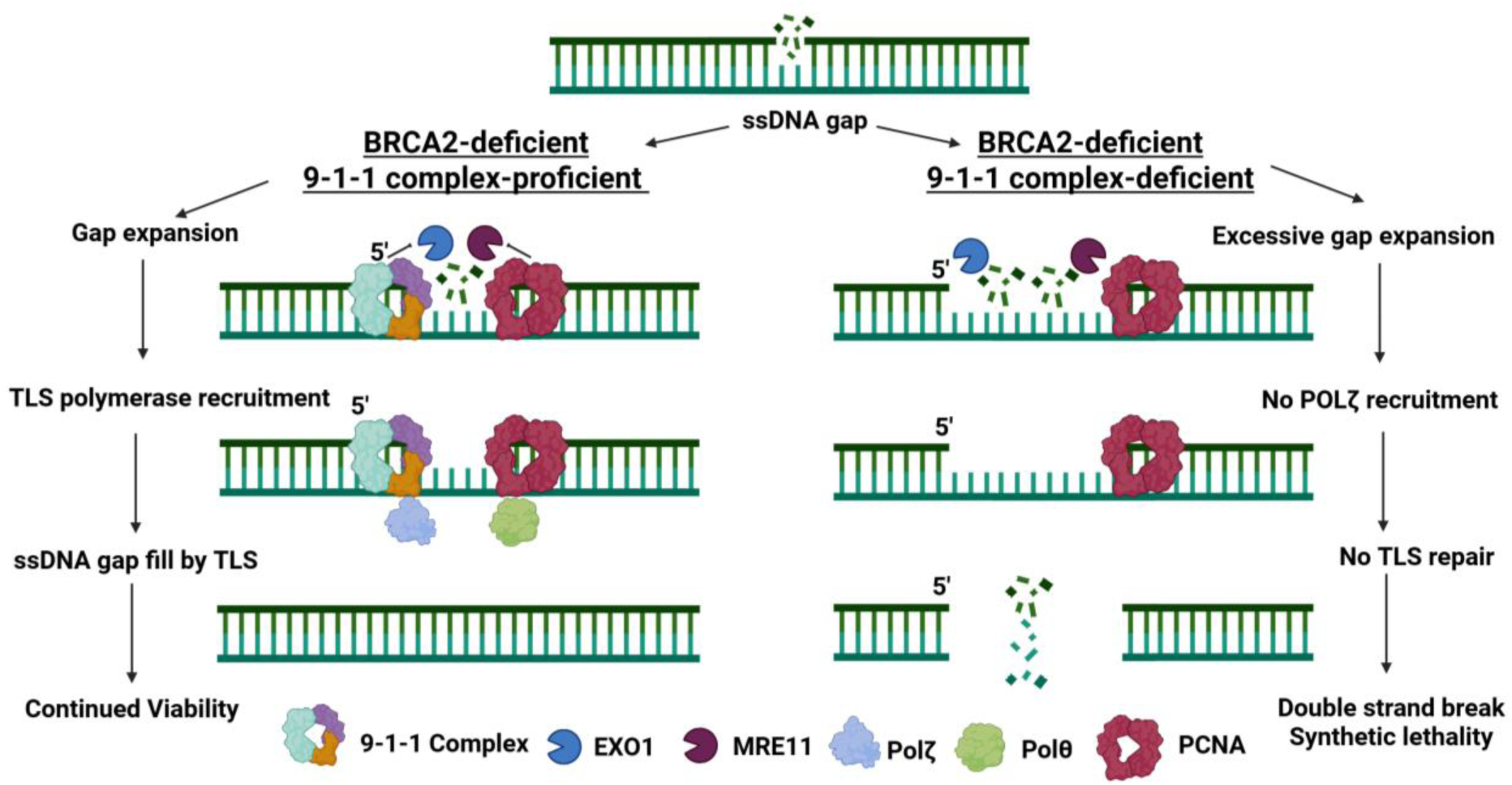
The 9-1-1 complex protects ssDNA gaps in BRCA2-deficient cancer. Working model of the mechanism causing synthetic lethality between BRCA2-deficiency and 9-1-1 complex loss. In BRCA2-deficient cells, the 9-1-1 clamp loads at the 5’ junction of PRIMPOL-generated ssDNA gaps, where it functions as a protective scaffold: it sterically hinders EXO1-mediated degradation to limit gap expansion and simultaneously recruits POLζ to promote gap filling via translesion synthesis. Loss of 9-1-1 disrupts this coordination, leading to pathological gap extension, replication catastrophe, and synthetic lethality.

### Therapeutic Implications

From a therapeutic standpoint, our findings position the 9-1-1 complex as a compelling target in BRCA2-deficient cancers. Unlike PARP inhibitors, which act by trapping repair intermediates, disrupting 9-1-1 function appears to destabilize ssDNA gaps. Because many BRCA2-mutant tumors eventually acquire PARPi resistance through restoration of fork stability or HR, inhibition of 9-1-1 could represent an orthogonal strategy that bypasses such resistance mechanisms.

The feasibility of drugging 9-1-1 directly remains an open question, but recent advances in targeting PCNA and TLS polymerases suggest that sliding clamps are tractable. Moreover, the clamp-loader RAD17 or its interactions with RFC subunits could offer alternative druggable interfaces. Small-molecule inhibitors that destabilize the complex or hinder the 9-1-1 complex loading onto DNA gaps at the 5’-end could selectively compromise the viability of BRCA2-deficient cells. Notably, the rescue observed upon EXO1 depletion suggests that combining 9-1-1 inhibition with modulators of nuclease activity or TLS could enhance the precision of replication stress-based therapies. Importantly, loss of the 9-1-1 complex was reasonably tolerated in BRCA2-proficient cell lines (DLD1, H1299, and RPE1), supporting a potential therapeutic window. However, the associated increase in DNA damage markers underscores the need for careful monitoring of toxicity in the development of 9-1-1-targeted strategies.

### Summary

In conclusion, our study defines a novel, ATR-independent role for the 9-1-1 complex in the protection and repair of ssDNA gaps, thereby positioning it as a central node in the survival network of BRCA2-deficient cells. Importantly, the selective synthetic lethality we observe in BRCA2-but not BRCA1-deficient contexts underscore the nuanced differences in how HR factors govern gap metabolism, with implications for tailoring therapeutic approaches to specific mutational backgrounds. By delineating this mechanism that prevents ssDNA gaps from becoming lethal, we have opened new avenues for synthetic lethality-based therapeutic strategies targeting the unique vulnerabilities of BRCA2-deficient tumors.

### Limitations of the study

This study primarily relies on isogeneic cell line pairs as models, which may not fully translate into *in vivo* systems or reflect the complexity of human tumors, limiting the ability to assess the physiological relevance, toxicity and therapeutic potential of targeting the 9-1-1 complex in BRCA2-deficient cancers. In this study, we predominantly focused on RAD9A and RAD1 as we were unable to generate a HUS1 knockout cell line model. HUS1 has been shown to function independently of the 9-1-1 complex through checkpoint signaling-independent roles and interactions with other proteins ^36,52,53^. Whilst its role in BRCA2-deficient cells has not been fully elucidated here, we have confirmed its loss to be synthetic lethal (Figure 1), suggesting a significant functional role as part of the 9-1-1 complex. Furthermore, while this study demonstrates that the 9-1-1 complex protects against ssDNA gap expansion and genomic instability in BRCA2-deficient cells, the precise molecular mechanism underlying double-strand break formation remains unresolved and is a subject of further work.

## Acknowledgments

We are indebted to The Genome Editing and Screening Core Facility for help with the CRISPR knockout screen. We thank Agnel Sfeir for their comments on the manuscript. This work was supported in part by MSK SPORE in Breast Cancer (CA247749) and CA286111. Figures were created with the aid of BioRender.

## Author Contributions

Conceptualization, S.N.P., H.E.G., J.S., and M.O; Methodology, S.N.P., H.E.G., and M.O; Validation, H.E.G., K.C., S.C., N.M., A.S., J.B., and M.O; Formal Analysis, H.E.G., K.C., S.C., N.M., A.S., J.B., J.S., and M.O; Investigation, H.E.G., K.C., S.C., N.M., A.S., J.B., J.S., and M.O.; Resources, S.N.P.; Writing – Original Draft, H.E.G., S.N.P.; Writing – Review & Editing, H.E.G., S.N.P.; Visualization, H.E.G., and N.M.; Supervision, S.N.P.; Project Administration, S.N.P.; Funding Acquisition, S.N.P.

## Inclusion and diversity statement

We support inclusive, diverse and equitable conduct of research.

### Cell culture and transfection conditions

The DLD1 WT, DLD1 BRCA2^-/-^, DLD1 RAD9A^-/-^, DLD1 RAD1^-/-^ H1299 WT, H1299 2D7 cell lines were grown in RPMI supplemented with 10% bovine growth serum, 10mM HEPES, 2mM L-Glutamine, 1mM sodium pyruvate, 100 I.U./ml Penicillin, and 100 μg/ml Streptomycin RPE1 WT, RPE1 BRCA1^-/-^ and RPE1 BRCA2 ^-/-^ cell lines were grown in DMEM HG supplemented with 10 % bovine growth serum, 100 I.U./ml Penicillin, and 100 μg/ml Streptomycin. Cells were grown at 37 **°**C with 5% CO_2_. DLD1 RAD9A^-/-^ and DLD1 RAD1^-/-^ were generated using the CRISPR-Cas9n system. Knockouts were verified through PCR and immunoprecipitation/western blot. Transfections with indicated siRNAs were performed using Lipofectamine RNAiMAX (Thermo Fisher Scientific) according to the manufacturers protocol. Cells transfected with siRNAs were used for experiments 48 hours after transfection.

### Whole genome CRISPR cas9 sequencing

Genome-wide CRISPR screen was performed using the human Brunello knockout (KO) library (targeting 19,114 genes with a total of 77,441 sgRNAs (4 sgRNAs per gene). Lentivirus carrying human CRISPR Brunello lentiviral pooled sgRNA library was produced in 293T cells. DLD1 isogenic cells were transduced (0.3 MOI) with the lentiviral Brunello sgRNA library to maintain >500x gRNA representation. Following puromycin selection (1.5 μg/mL) surviving cells were allowed to proliferate for 14 days, with cell pellets harvested in triplicate at Day 0 and Day 14. Guide RNA cassettes were amplified from extracted genomic DNA to generate Illumina sequencing libraries. Namely, 3 µg of genomic DNA was added per 50 µl PCR reaction using staggered primers to increase base diversity. PCR products were then pooled and purified using QIAquick PCR purification kits (Qiagen). Pooled samples were sequenced by Illumina Novaseq. Data was analyzed using the Mageck-vispr workflow and beta values were calculated using MAGeCK MLE (maximum likelihood estimation) which represent the estimated effect size of gene perturbation.

### Clonogenic assay

Cells were treated as stated, trypsinized and replated in 10-cm dishes at appropriate concentrations in triplicates. Fourteen-twenty days after replating, cells were washed with PBS, fixed with methanol, and stained with crystal violet. Colonies with more than 50 cells were counted. Colony counts were normalized to control cells of the same genotype. Plating efficiency was calculated as number of colonies formed/ number of cells seeded.

### Western blot

Cells were harvested by scraping, washed with PBS, and pelleted. Cells were lysed in RIPA buffer (25 mM Tris-HCl pH 7.6, 150 mM NaCl, 0.1% SDS, 1% NP-40, 1% sodium deoxycholate) with 1 mmol/L protease inhibitor PSMF and HALT protease inhibitor cocktail (Thermo Scientific). Protein samples were sonicated and centrifuged to obtain protein lysate. The concentration of protein lysate was quantified using the BCA kit per manufacturer’s instructions (Thermo Scientific). Protein lysates were normalized and diluted in SDS-PAGE reducing sample buffer (Thermo Scientific) and reducing agent (Thermo Scientific) before denatured at 10 minutes at 95°C. The samples were loaded into Bis-Tris or Tris-Acetate precast gels (Invitrogen) for SDS-gel electrophoresis and transferred onto Immobilon-PVDF membrane (Millipore). Membranes were blocked for 1 hour at room temperature with 5% in TBS and incubated with primary antibody for either 2 hours at room temperature or overnight at 4 °C. Membranes were washed with PBS + 0.1% Tween 20 (PBST), incubated with secondary antibody for 1 hour at room temperature, and washed with PBST. Proteins were then detected with chemiluminescence and developed on film.

### Cell cycle analysis

Cell cycle analysis was performed with Click-iT™ EdU Cell Proliferation Kit for Imaging, Alexa Fluor™ 488 dye (Thermo Fisher Scientific) according to the manufacturer’s instructions.

### DNA fiber assay

To measure replication rate, cells were pulse-labeled with 20 µM 5-chloro-2’-deoxyuridine (CldU, Sigma-Aldrich) for 20 min, washed three times with PBS, and followed by a second label with 200 µM 5-iodo-2’-deoxyuridine (IdU, Sigma-Aldrich) for 1 hour and the cells collected. For DNA fiber assay treated with S1 nuclease, cells were incubated with 20 µM IdU for 1hr before being permeabilized with CSK100 (100mM NaCl, 10mM MOPS, pH 7, 3mM MgCl2, 300mM sucrose, 0.5% Triton X-100) for 10 min at room temperature. Cells were washed carefully with PBS, washed once with S1 nuclease buffer (30mM sodium acetate, 10mM zinc acetate, 5% glycerol, 50mMNaCl, pH 4.6), and treated with 20U/mL S1 nuclease suspended in S1 nuclease buffer for 30 min at 37 °C and then the cells were collected. For DNA fiber assay to measure post-replicative repair, cells were pulse labelled with 20 µM IdU for 30 minutes before washed with PBS and incubated with 200 ng/mL nocodazole for 14 hours. In the last 4 hours of nocodazole treatment cells are pulsed with 20 µM CldU, before the cells were collected. Additional drug treatments were conducted as stated.

All DNA fiber assays were spread and stained in the same way. Cells were collected in PBS-0.1% BSA with a cell scraper, centrifuged for 5 min at 4 **°**C, and re-suspended in PBS. 2 µl of cells were mixed with 8 µl of lysis buffer (200mM Tris–HCl, pH 7.5, 50mMEDTA, 0.5% SDS) on the top of a positively charged glass slide and incubated for 8 min at room temperature. Slides were tilted at a 30-45° angle to spread the fibers at a constant low speed. Slides were dried for 10-15 min at room temperature before being fixed in freshly prepared 3:1 glacial acetic acid for 15 min at-20 °C. Slides were then dried and stored at 4°C overnight. Slides were rehydrated in 2 x 5 min washes in PBS. DNA was denatured with 2.5 M HCL for 1 hour at room temperature. Slides were washed 3 x 5 min with PBS before being blocked for 45 min with warmed 5% BSA at 37 °C. Excess liquid was gently removed, and slides were incubated with 30 µl of primary antibody with a coverslip in a dark humid chamber for 1h 30 min. Slides were placed in PBS for 1-2 min to remove coverslips. Slides were washed with 0.05% Tween-20 in PBS for 3 x 5 min and incubated with 30 µl of secondary antibody with a coverslip in a dark humid chamber for 1h. Slides were placed in PBS for 1-2 min to remove coverslips. Slides were washed with 0.05% Tween-20 in PBS for 3 x 5 min and dried. A coverslip was mounted on the slides with Prolong Diamond Antifade Mountant (Thermofisher Scientific) and left to dry at room temperature in the dark. Slides were stored at 4°C.The following primary antibodies were used: mouse anti-BrdU (1:20) and rat anti-BrdU (1:100). The following secondary antibodies were used: anti-mouse IgG1 Alexa Fluor 546 and anti-rat Alexa Fluor 488. All antibodies were made in 1% BSA 0.05% Tween-20 in PBS. DNA fibers were visualized at 488 nm on a Nikon Confocal Microscope. At least 100 fibers per condition per replicate were quantified.

### Single Molecule Analysis of ssDNA gaps (SMAss)

Cells were transfected with siRNA for 48 hours. After 24 hours, cells were pulse labelled with 200 µM IdU for 24 hours. DNA combing was performed essentially as previously described, but under non-denaturing conditions ^54^. Cells were harvested by gentle trypsinization and counted before being resuspended in PBS at 1.5×10^5^ cells. Cells were prepared into 1.2% melting agarose plugs and underwent proteinase K treatment in ESP buffer (0.5M EDTA pH=8, 10%(v/v) Sarkosyl/0m5M EDTA pH=8.0, 20mg/ml Proteinase K) at 50°C for 16-18 hours. Plugs were washed with TE buffer (10mM Tris-HCL, 1mM EDTA pH=8) on a rotating mixer for 3x 1-hour washes and a final wash for 3.5 hours. Plugs are incubated in 0.5 MES at 68°C for 20 mins before incubation overnight at 42°C with β-Agarase. Samples are diluted in 1 mL 0.5M MES solution and poured into a reservoir before using the DNA combing machine to stretch DNA fibers on silanized coverslip at a constant rate of 300 µm/s. Coverslips are dehydrate at 65°C for 2 hours.

### Foci immunofluorescence

Cells were grown in Millicell EZ chamber slides (Merck Millipore) and fixed post-treatment. Cells were washed with PBS and simultaneously fixed and permeabilized with 4% PFA and 0.2% Triton-x100 in PBS at room temperature. Additional extraction with 0.5% Triton-x100 in PBS was done at room temperature for 10-15 minutes. Cells were treated with methanol for 30 secs before being washed with PBS. Cells were blocked in 3% BSA in PBS for 1 hour at 37 °C. Blocked cells were incubated for 4 hours with the primary antibody diluted in 3% BSA in PBS before being washed with PBS 0.1% triton. Cells were incubated with secondary antibody diluted in 3% BSA in PBS for 1 hour and then washed with PBS 0.1% triton. Wells were removed and dried with a vacuum and cover slips were mounted with Prolong Diamon Antifade Mountant with DAPI (Thermofisher Scientific). Slides were visualized at 488 nm on a Nikon Confocal Microscope. Foci of at least 100 cells per condition per replicate were quantified.

### Comet Assay

Cells were gently scraped and transferred to a centrifuge tube suspended in ice-cold 1X PBS. Cells were counted and transferred to a fresh centrifuge before centrifugation and resuspension at 5 x 0^5^ in ice-cold 1X PBS. Cell suspensions were combined with 1% low melting agarose (cooled in a water bath to 37⁰C) at a ratio of 1:10 (v/v) and 50 ul was spread onto a CometSlide (R&D Systems). Slides were placed flat at 4⁰C in the dark for 15-30 mins to allow for gelling time. Slides were immersed in 4⁰C Lysis Solution (R&D Systems) and incubated overnight at 4⁰C. Excess lysis buffer was drained from the slides, and then gently immersed in pre-cooled 1X Neutral Electrophoresis Buffer (10X: 1mM Tris Base, 3mM Sodium Acetate adjusted to pH 9.0 with glacial acetic acid) at 4⁰C for 30 minutes before being placed in an electrophoresis tank. The electrophoresis tank was filled with 1x Neutral Electrophoresis buffer (pre-cooled at 4⁰C) to a level no higher than 0.5 cm above the slides. The power supply was set to a voltage where 1 volt per cm measured electrode to electrode. and ran for 45 mins at 4⁰C. Excell Neutral electrophoresis buffer was drained before the slides were immersed in DNA precipitation Solution (1 M NH4Ac, 95% EtOH) for 30 minutes at room temperature. Slides were then immersed in 70% EtOH for 30 minutes at room temperature. Slides were dried at 37⁰C for 10-15 mins. 100 µl of SYBR Gold (diluted in TE buffer, 7.5 pH) was placed on each circle of drained agarose and stained for 30 minutes at room temperature in the dark. Slides were immersed in ice-cold water for 30 minutes and then dried completely at 37⁰C. Slides were visualized at 488 nm on a Nikon Confocal Microscope. At least 50 comets were measured per condition. The tail moment was calculated as a percentage of the sum intensity of the comet tail over the sum intensity of the whole comet.

## Data analysis and Statistics

Statistical analysis for the various experiments was performed using GraphPad Prism. Results are presented as mean ± standard error of the mean (SEM). A p-value of <0.05 by Student’s t-test was considered statistically significant. ns: non-significant, * Indicates p<0.05, **: P<0.005, ***: P<0.001.

## Supplementary Figures and Tables

**Supplementary Figure 1:**
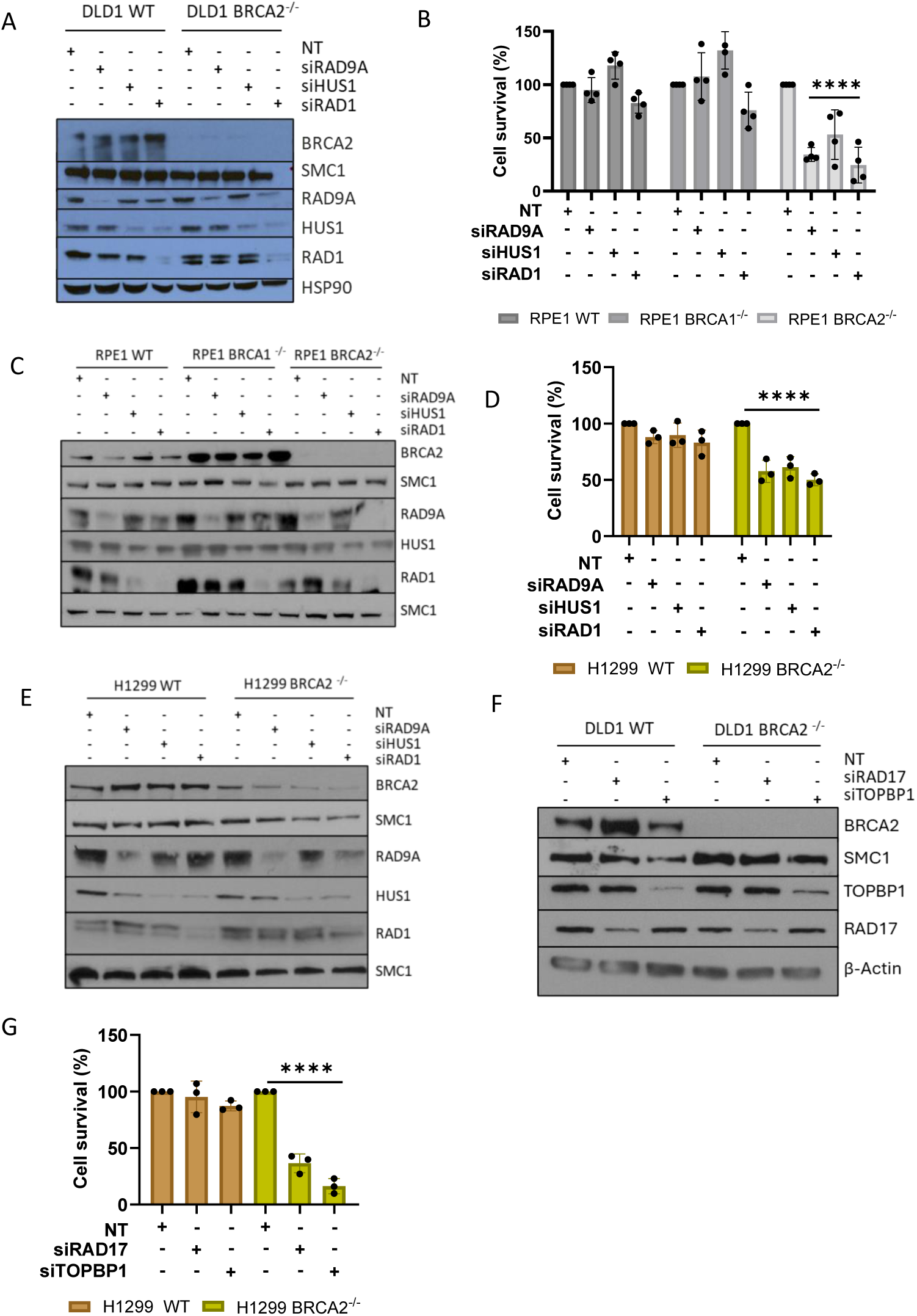
Loss of BRCA2 and 9-1-1 complex is synthetic lethal across multiple cell line models. (A) Western blot analysis of siRNA knockdown for clonogenic assay in Figure 1B. (B) Clonogenic survival assay in the BRCA2/BRCA1-proficient and-deficient RPE1 cells with siRNA depletion of RAD9A, HUS1 and RAD1, n = ≥ 3, mean + SEM, ns= p>0.05, ****p <0.0001, paired t test. (C) Western blot analysis of siRNA knockdown for (B). (D) Clonogenic survival assay in the BRCA2-proficient and-deficient H1299 cells with siRNA depletion of RAD9A, HUS1 and RAD1, n = ≥ 3, mean + SEM, ns= p>0.05, ****p <0.0001, paired t test. (E) Western blot analysis for (D). (F) Western blot analysis of siRNA knockdown for clonogenic assay in Figure 1C-1D. (G) Clonogenic survival assay in the BRCA2-proficient and-deficient H1299 cells with siRNA depletion of RAD17 and TOPBP1, n = 3, mean + SEM, ns= p>0.05, ****p <0.0001, paired t test.

**Supplementary Figure 2:**
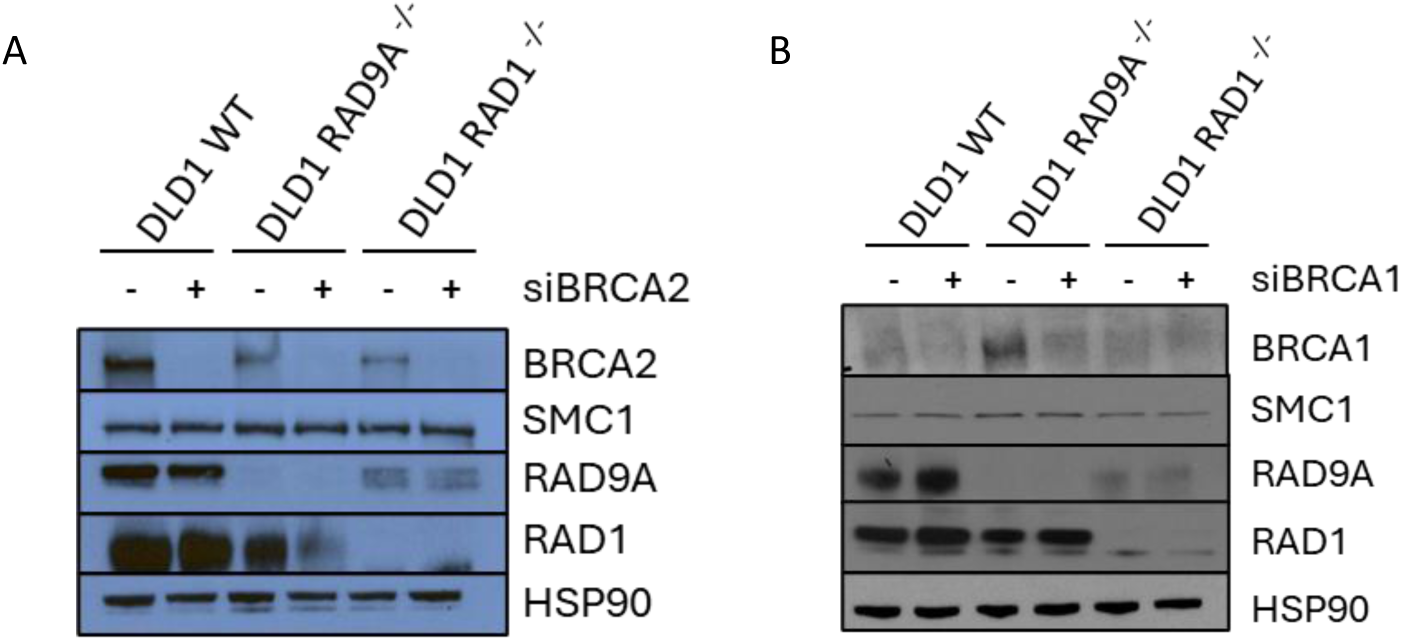
RAD9A and RAD1-deficient cells demonstrate synthetic lethality with loss of BRCA2 but not BRCA1. (A) Western blot analysis of siRNA knockdown for clonogenic assay in Figure 1E. (B) Western blot analysis of siRNA knockdown for clonogenic assay in Figure 1F.

**Supplementary Figure 3:**
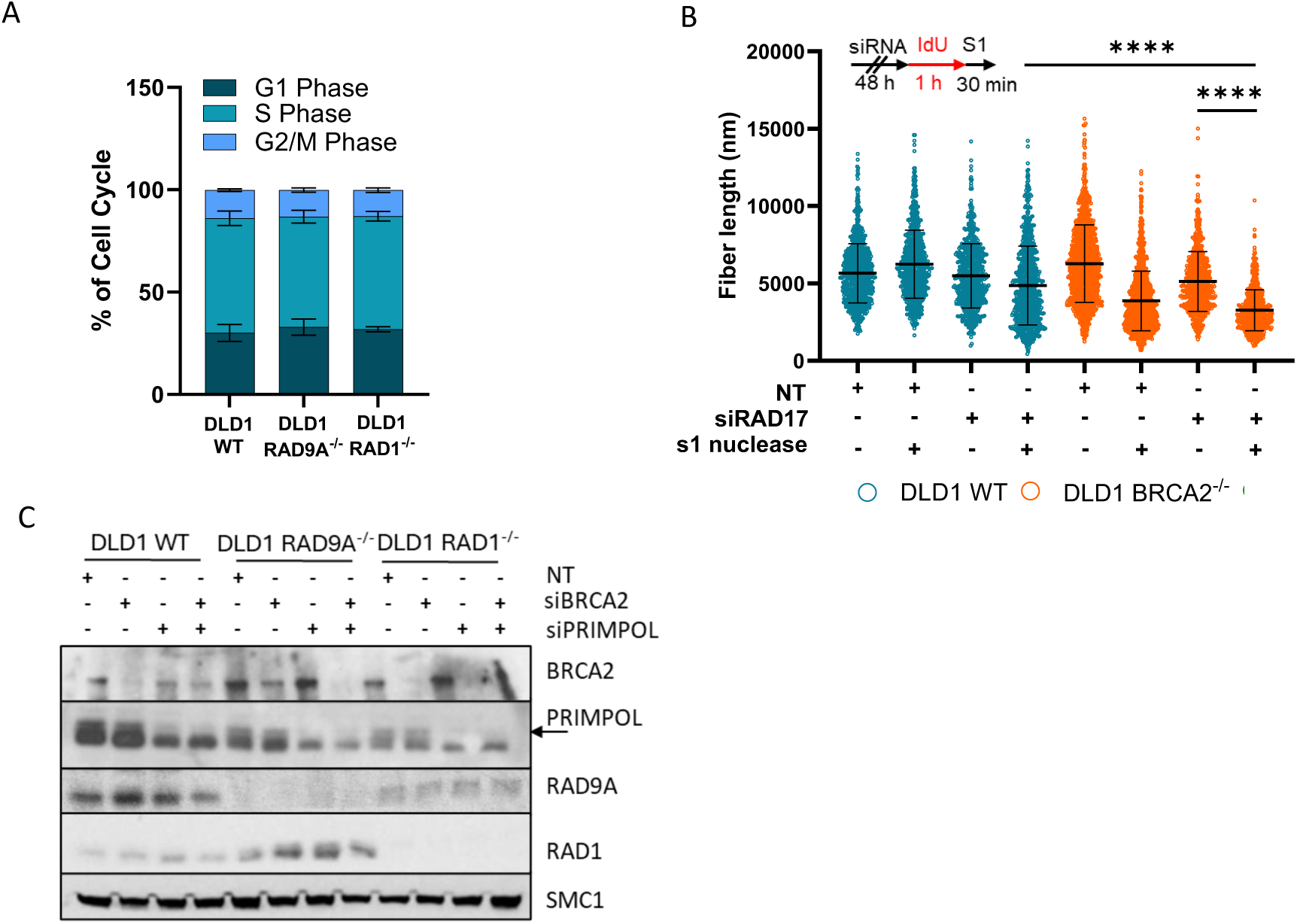
Loading of the 9-1-1 complex is required to prevent PRIMPOL-mediated ssDNA gap accumulation in BRCA2-deficient cells. (A) Cell cycle analysis of RAD9A/RAD1-deficient and proficient cell lines, n=3, mean + SEM. (B) Analysis of DNA fiber length after incubation with and without S1 nuclease. Set up of experiment (top) and dot-plot (bottom). Depletion of RAD17 in BRCA2-deficient and proficient cells, n = ≥ 3, at least 100 fibers were analyzed per condition, per replicate, mean + SEM, ****p <0.0001, paired t test. (C) Western blot analysis of PRIMPOL depletion in RAD9A/ RAD1-deficient and proficient cells.

**Supplementary Figure 4:**
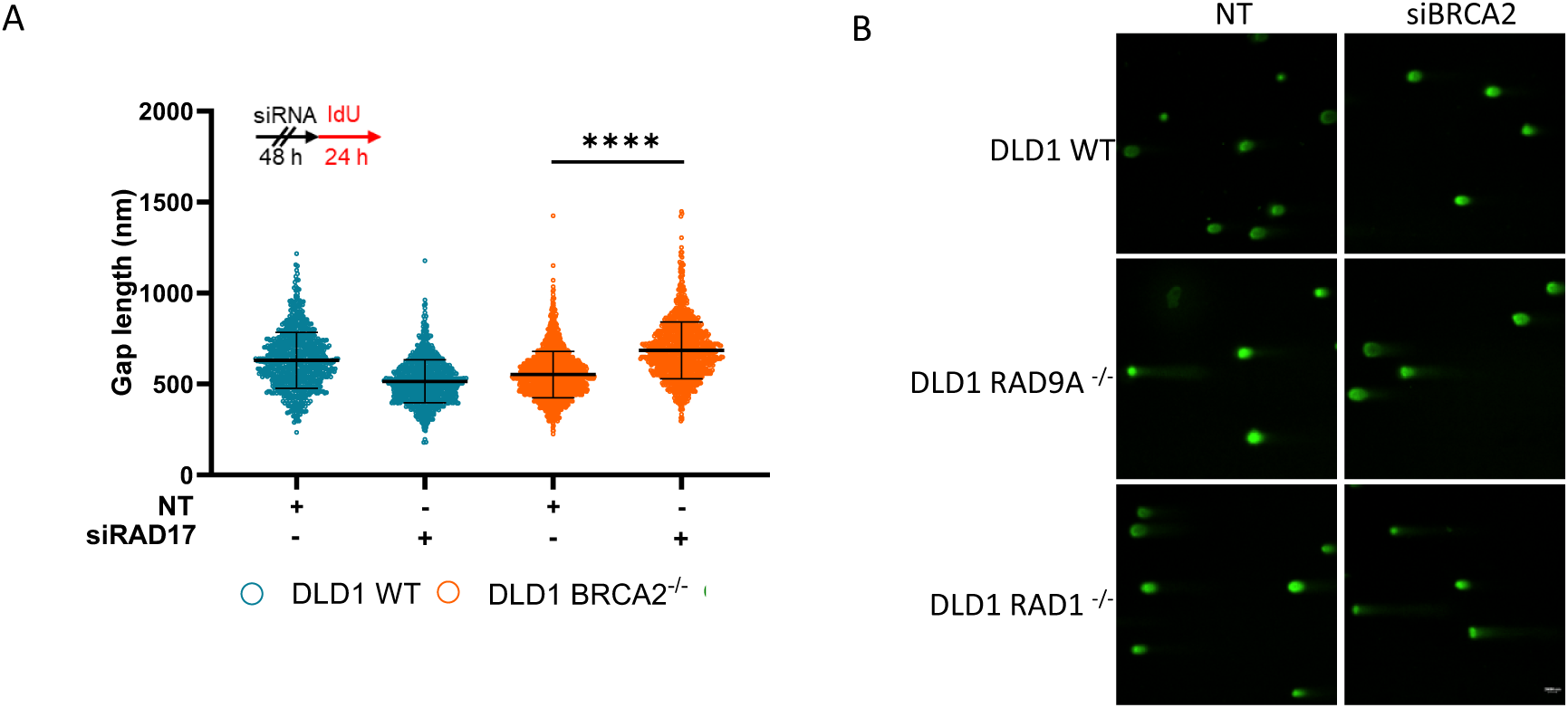
Loading of the 9-1-1 complex is required to prevent ssDNA gap lengthening and genomic instability in BRCA2-deficient cells. (A) Single Molecule analysis of ssDNA gaps (SMass). Schematic of experiment (top) dot-plot (bottom) with depletion of RAD17 in BRCA2-deficient and proficient cells, n = 3, at least 100 ssDNA gaps were analyzed per condition, per replicate, mean + SEM, ****p <0.0001, paired t test. (B) Example images of comet assay in Figure 3H.

**Supplementary Figure 5:**
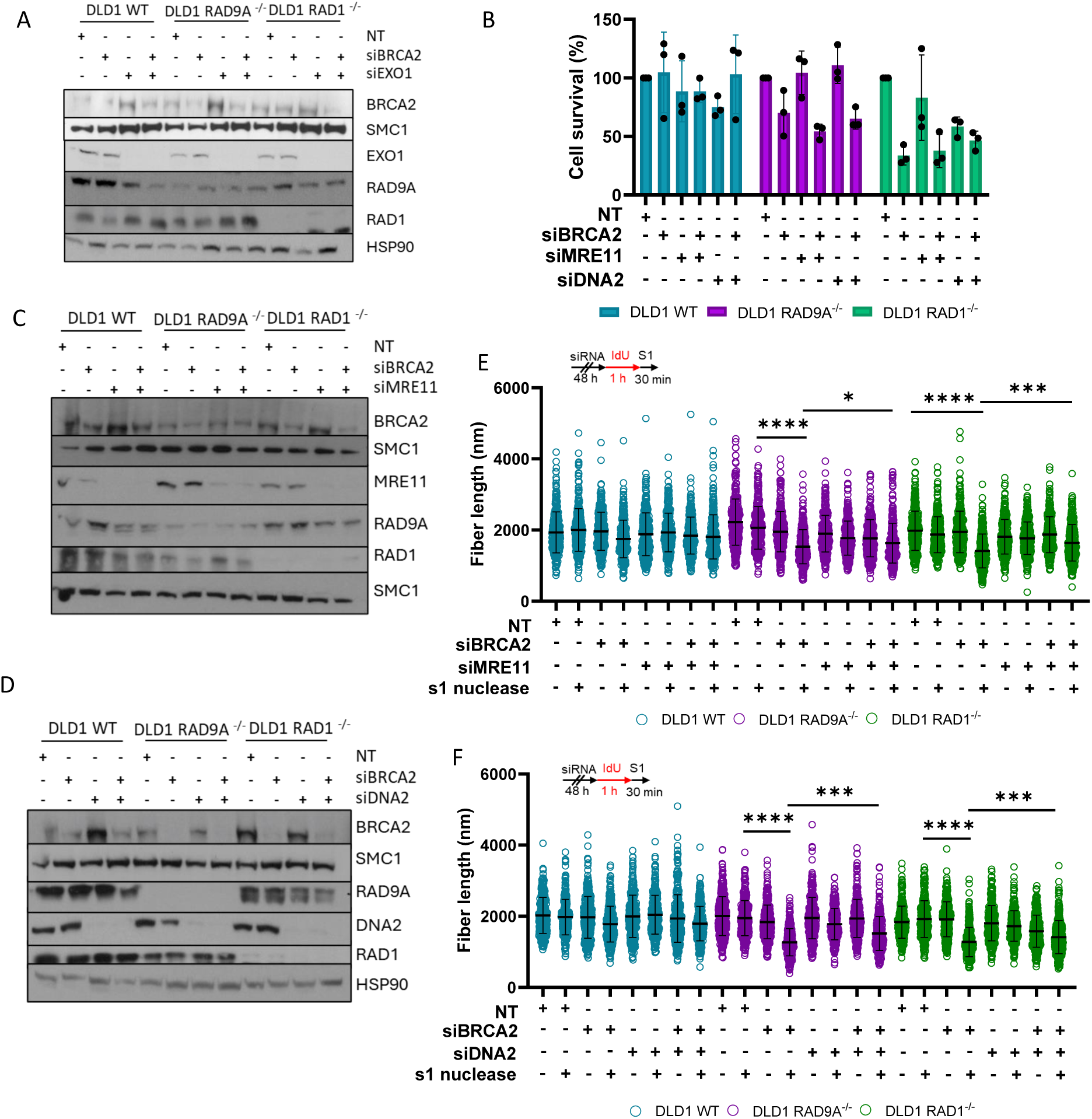
The 9-1-1 complex protects ssDNA gaps from EXO1-directed expansion. (A) Western blot analysis of siRNA knockdown in Figure 4A. (B) Clonogenic survival assay in RAD9A/ RAD1-deficient and proficient cells with siRNA depletion of BRCA2, MRE11 and/or DNA2, n = ≥ 3, mean + SEM, ns= p>0.05, ****p <0.0001, paired t test. (C-D) Western blot analysis of siRNA knockdown for (B). (E-F) Analysis of DNA fiber length after incubation with and without S1 nuclease. Set up of experiment (top) and dot-plot (bottom). Depletion of BRCA2 with/without (E) MRE11 (F) DNA2 depletion, n = ≥ 3, at least 100 fibers were analyzed per condition, per replicate, mean + SEM, ns= p>0.05, ****p <0.0001, paired t test.

**Supplementary Figure 6:**
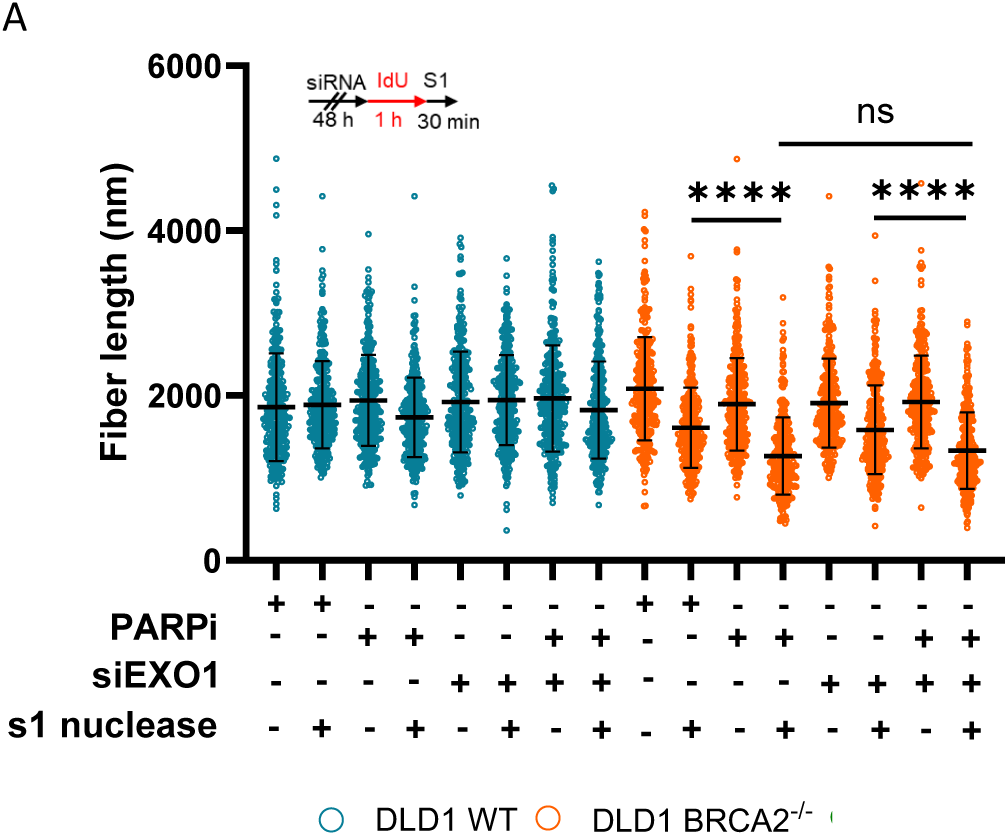
EXO1 depletion does not rescue Olaparib-induced ssDNA gaps. (A) Analysis of DNA fiber length after incubation with and without S1 nuclease. Set up of experiment (top) and dot-plot (bottom). Depletion of EXO1 with and without Olaparib treatment in BRCA2-proficient and deficient cells, n = ≥ 3, at least 100 fibers were analyzed per condition, per replicate, mean + SEM, ns= p>0.05, ****p <0.0001, paired t test.

**Supplementary Figure 7:**
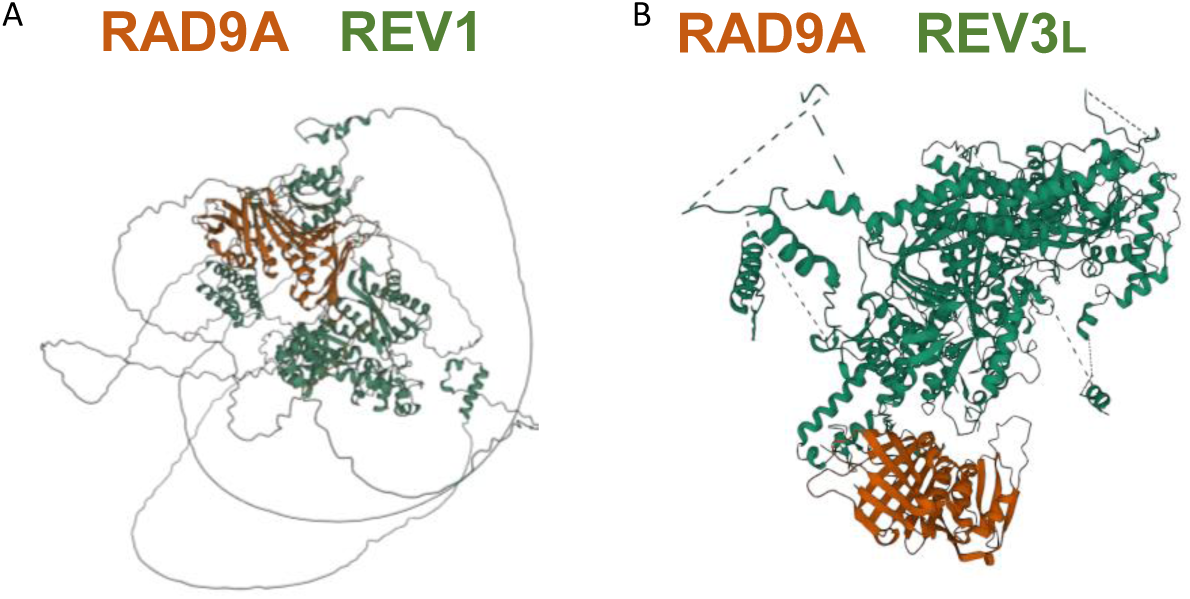
RAD9A is predicted to interact with REV1 and REV3L. Predicted structural interactions between RAD9A and (A) REV1 and (B) REV3L generated through Predictomes dataset. Orange is RAD9A structure and green is REV1 and REV3L structures.

**Supplementary Figure 8:**
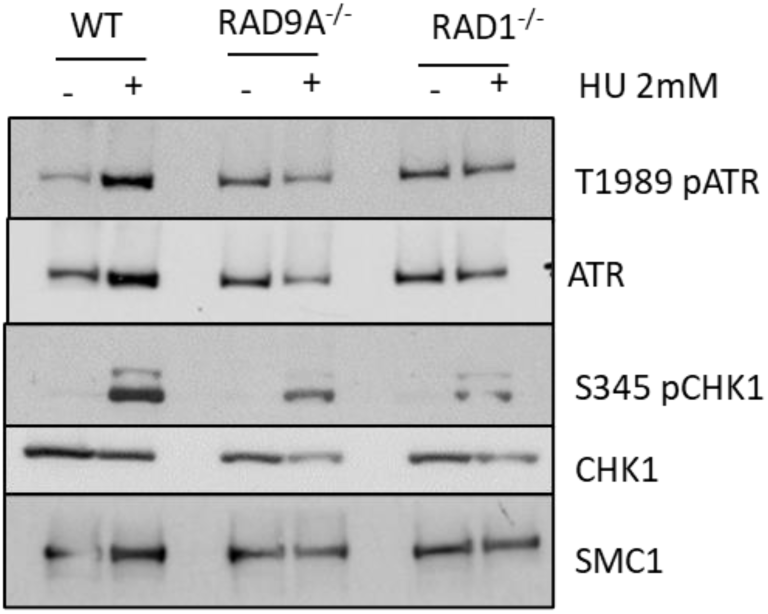
Loss of the 9-1-1 complex reduces ATR signaling. (A) Western blot analysis of ATR and CHK1 signaling when RAD9A/RAD1-deficient and proficient cells are treated with 2mM hydroxyurea.

